# Emergence of disease-specific endothelial and stromal cell populations responsible for arterial remodeling during development of pulmonary arterial hypertension

**DOI:** 10.1101/2023.09.06.555321

**Authors:** Nicholas D Cober, Emma McCourt, Rafael Soares Godoy, Yupu Deng, Ken Schlosser, Anu Situ, David P Cook, Sarah-Eve Lemay, Timothy Klouda, Ke Yuan, Sébastien Bonnet, Duncan J Stewart

## Abstract

Pulmonary arterial hypertension (PAH) is a severe and lethal pulmonary vascular disease characterized by arteriolar pruning and occlusive vascular remodeling leading to increased pulmonary vascular resistance and eventually right heart failure. While endothelial cell (EC) injury and apoptosis are known triggers for this disease, the mechanisms by which they lead to complex arterial remodeling remain obscure. We employed multiplexed single-cell RNA sequencing (scRNA-seq) at multiple timepoints during the onset and progression of disease in a model of severe PAH to identify mechanisms involved in the development of occlusive arterial lesions. There was significant loss of arterial volume as early as 1-week by microCT, preceding any evidence of occlusive arteriopathy, consistent with early arteriolar dropout. Maximal arterial pruning was seen by 5 to 8 weeks, with signs of progressive occlusive remodeling. Analysis of the scRNA-seq data resolved 44 lung cell populations, with widespread early transcriptomic changes at 1 week affecting endothelial, stromal and immune cell populations. Notably, this included emergence of a relatively dedifferentiated (dD) EC population that was enriched for *Cd74* expression compared to general capillary (gCap) ECs which were primed to undergo endothelial-mesenchymal transition, as evidenced by RNA velocity analysis. However, at late timepoints (5 and 8 weeks), activated arterial ECs (aAECs) were the only cell population exhibiting persistent differential gene expression. This was characterized by a growth regulated state, including high expression of *Tm4sf1*, a gene implicated in cancer cell growth, which was also expressed by a smooth muscle (SM)-like pericyte cluster. Both these populations were localized to regions of arterial remodeling in the rat model and PAH patients, with aAECs contributing to intimal occlusive lesions and SM-like pericytes forming bands of medial muscularization. Together these findings implicate disease-specific vascular cells in PAH progression and suggest that TM4SF1 may be a novel therapeutic target for arterial remodeling.

## Introduction

Pulmonary arterial hypertension (PAH) is a severe and lethal disease defined hemodynamically by precapillary arterial remodeling.^1^ While the pathobiology of PAH is incompletely understood, endothelial cell (EC) dysfunction has long been known to contribute to increased vascular tone and remodeling.^2^ More recently, EC injury and apoptosis has been recognized as a central trigger for initiation of disease,^3^ and underlying mutations in BMPR2, which represent the most common cause of hereditary PAH,^4^ have been shown to increase susceptibility of ECs to injury and apoptosis.^5^ However, it is still uncertain how EC apoptosis leads to loss of vasculature and the development of complex arterial remodeling and plexiform lesions, which are hallmark features of advanced PAH.^3^

Recently, it has been reported that perturbed pulmonary hemodynamics is necessary for the development of occlusive arterial remodeling,^6^ possibly as a result of increased intimal shear stress within the lung arteriolar bed which could contribute to ongoing endothelial injury. We have proposed that there are at least two distinct phases during the development of PAH.^3^ The first represents the initial response to endothelial injury and apoptosis, which leads directly to loss of fragile precapillary arterioles and progressive arteriolar pruning as the initial cause of perturbed lung hemodynamic forces that drives complex arterial remodeling. At later stages of disease, ongoing hemodynamic abnormalities likely result in persistent arteriolar EC injury, perpetuating arterial remodeling and disease progression. Single-cell transcriptomics is a powerful tool that can elucidate the transcriptional changes underlying the onset and progression of disease at the resolution of individual cells, thereby providing new insights into the mechanisms driving the functional and structural vascular abnormalities of PAH. Single-cell RNA sequencing (scRNA-seq) has been widely employed to better characterize lung cell populations under homeostatic conditions,^7,8^ and in various disease states.^9,10^ Recently, several studies have employed scRNA-seq to characterize transcriptomic profiles in experimental models and human disease.^11–14^ However, all of these studies have primarily focused on the late stages of disease, whereas the mechanisms leading to the development of PAH in the early stages may be quite distinct. Multiplexed scRNA-seq analysis can increase the number of samples analyzed at one time, making it feasible to study multiple timepoints with increased biological replicates by reducing costs.^15^

Therefore, we employed multiplexed scRNA-seq to map out the temporal changes in transcriptomic profiles and cell populations during onset and progression of PAH in the SU5416-chronic hypoxia (SU/CH) rat model, focusing mainly on the vascular cell populations. We identified two main phases of disease characterized in the early stages (i.e., 1 week post SU) by widespread changes in endothelial, stromal, and immune cell populations, and the emergence of a de-differentiated (dD) EC cluster, which was primed to undergo endothelial to mesenchymal transition (EndMT) as was seen by RNA velocity analysis. At later stages of disease (i.e., 5 and 8 weeks), there was a surprising normalization of global transcriptional activity in most lung cell populations, with the notable exception of the activated arterial ECs (aAECs) cluster, which continued to exhibit persistent and robust changes in gene expression. Interestingly, these cells were highly localized to regions of complex arteriolar remodeling, together with the smooth muscle (SM)-like pericytes, both of which expressed *Tm4sf1*, a marker previously implicated in cancer biology.

## Methods

### Sugen5416 – chronic hypoxia model of PAH

All animal experiments were approved by the University of Ottawa Animal Care Committee and conducted according to the guidelines from the Canadian Council for Animal Care. Male Sprague Dawley (SD) rats (6-10 weeks old) were administered with a single dose of Sugen 5416 (SU) (20mg/kg, Tocris) delivered in CMC vehicle (as previously described)^16^ or DMSO and placed in hypoxia chambers (10% O2, Biospherix) for three weeks, followed by 5 weeks of normoxia. Animals at baseline, 1, 3, 5, and 8-weeks (**Figure 1A**) were anaesthetized by an i.p. injection of ketamine (100 mg/kg) and xyalzine (10 mg/kg) and right ventricular systolic pressure (RVSP) was measured by a blinded analyst, as previously described.^17^ Lungs were either isolated for tissue digestion (described below) or formalin fixed and paraffin embedded for subsequent histological analysis. The heart was excised and right ventricular hypertrophy (RVH) was assessed by the ratio of right ventricle (RV) to the left ventricle plus septum (LV+S), as previously described.^17^

**Figure 1:**
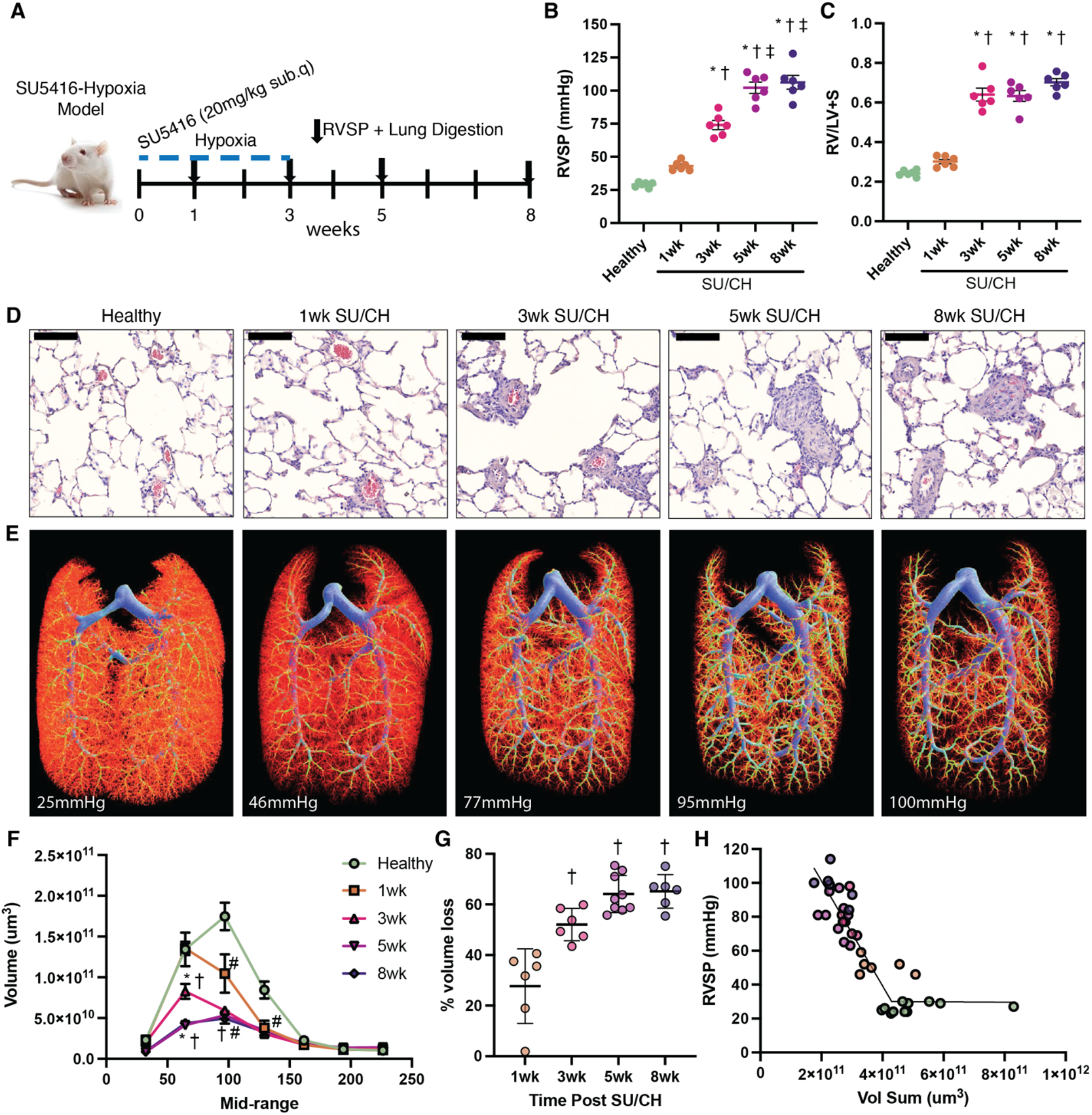
Characterization of Sugen-Chronic Hypoxia (SU/CH) model of PAH. A) Timeline of the SU/CH model and relevant end of study sampling. B) Right ventricular systolic pressure (RVSP). C) Right ventricular hypertrophy (RV/LV+S). D) Representative H&E histological images showing progression of arterial remodeling associated during development of pulmonary hypertension (PH). E) Representative image of micro computed tomography (MicroCT) of lung arterial angiograms during the development of PH. F) Summary of quantitative analysis of arterial volumes by MicroCT showing the greatest reduction in arterioles under 150µm in diameter. G) Percent volume loss relative to healthy control in arteries under 200µm in diameter. H) Relationship between arterial volume (<500µm diameter) and RVSP during PH development. Data represented as mean ± SEM, n = 6 – 12. * p < 0.05 for 3-, 5-, 8- weeks SU/CH vs control, † p < 0.05 for 3-, 5-, 8- weeks vs 1-week, ‡ p < 0.05 for 5-, 8- weeks vs 3-weeks, # p < 0.05 for 0.01 for all timepoints vs control. Scale bar represents 100μm.

### Sample preparation for micro-computed tomography (microCT)

Lungs were prepared for microCT, as previously described.^18^ In brief, animals were thoroughly perfused with heparinized saline (LEO Pharma Inc.) through the jugular vein with a closed chest and mechanical ventilation, using pressure matching the measured RVSP. Subsequently, chests were opened, and an additional perfusion of heparinized saline cleared residual blood from the lungs. Barium – gelatin mix (E-Z-EM Canada Inc; Sigma-Aldrich) was perfused until consistent pressure matching RVSP was obtained. Lungs were cooled to 4°C and inflated with formalin through the trachea. Lung samples were stored at 4°C in formalin for 2 days, washed with PBS, and stored in ethanol until images were acquired with desktop microCT (SkyScan 1272, Bruker microCT). Image analysis was performed with CTAn software (Bruker microCT), and CTVox software (Bruker microCT).

### H&E staining and histology

Hematoxylin and eosin (H&E) staining was performed on paraffin sections. Sections (5µm thick) were deparaffinized and rehydrated as previously described using a series of xylene, ethanol, and water washes. Sections were stained with hematoxylin for up to 8 min and washed with water. Then incubated in Scott’s solution for 2 min, washed and soaked in eosin stain for 3 min. Slides were washed in water, and then dehydrated in a reverse series of ethanol and xylene rinses. Slides were mounted with Cytoseal XYL media (Thomas Scientific) and left to dry overnight. Slides were scanned with Panoramic Desk (3D Histech, Budapest, Hungary) and analyzed with CaseViewer (3D Histech).

### Lung digestion and single cell preparation

Lung digestion was performed in a similar manner as previously described.^19^ In brief, animals were perfused with 20ml of 0.9% saline containing 25U/ml heparin (LEO Pharm Inc.) through the right ventricle, to drain the blood from the lungs. Lung tissue was isolated, minced with scissors and placed in Hank’s Balanced Salt Solution (HBSS, Gibco) on ice. Enzymatic digestion was performed with 2500U Collagenase (Worthington), 30U Neutral Protease (Worthington), and 165U DNase I (Sigma Aldrich) in HBSS, samples were loaded into gentleMACS C tubes (Milltenyi Biotec), and placed on the OctoMACS dissociators (Milltenyi Biotec) using program 37C_LDK_m_1 for ∼30min. Samples were immediately placed on ice, filtered through 70µm cell strainers (Falcon), and washed with phosphate buffered saline (PBS, Gibco) with 2mM EDTA (ThermoFischer Scientific) and 0.5% bovine serum albumin (BSA, Wisent) (PEB). Dissociated cell samples were spun at 400g for 5min at 4°C and washed again. Samples were incubated in 1x red blood cell lysis (eBioscience) for 3min, diluted with PEB, and filtered again through 70µm cell strainers (Milltenyi Biotec). Cells were washed, as above and counted with Trypan Blue (ThermoFisher Scientific) on a Cell Drop BF (DeNovix).

### Multiplex single cell sample processing

Individual biological samples were barcoded using the MULTI-seq protocol described by McGinnis et al.^15^ In brief, single cells were incubated with 1:1 molar ratio of anchor: unique barcode oligonucleotide sequence for 10 min at room temperature with gentle mixing. Followed by addition of the co-anchor to stabilize barcodes within membranes which was incubated for 5min on ice. Barcoded cells were washed in PBS, counted as above, and pooled at an equal ratio of cells per biological replicates. Samples with viability > 80% were sent for further processing. Pooled samples were processed using 10x Genomics Chromium (10x Genomics).

### Processing of single cell RNA sequencing libraries

RNA library construction was performed using 10x Genomics Single Cell 3’ RNA sequencing kit as previously described.^19^ This experiment was performed with unique biological replicates on different dates, and data was later merged. Libraries were created and sequenced with NextSeq500 (Illumina) for an estimated 37554 cells with 21879 mean reads per cell, 2372 median UMI per cell, and 1118 median genes per cell (first experiment) and 35546 cells with 22322 mean reads per cell, 3152 median UMI per cell, and 1370 median genes per cell (experiment repeat). CellRanger (10x Genomics, version 6.1.2) was used to process raw sequencing reads using the Ensembl104 rat transcriptome annotations with additional inclusion of Pecam.

### Quality control and single cell data analysis

Filtered feature barcode matrices were imported into R package Seurat (https://github.com/satijalab/seurat) for subsequent analysis.^20^ Quality control was performed within Seurat to remove cells with >30% mitochondrial transcripts and cells with low complexity (<200 detected genes). Barcodes were demultiplexed using a manual threshold method outlined within the deMULTIplex R package (https://github.com/chris-mcginnis-ucsf/MULTI-seq).^15^ Cells with multiple barcodes (doublets) or with no identifiable barcodes were removed from analysis. An additional round of doublet removal was performed with scDblFinder (https://github.com/plger/scDblFinder).^21^ A total of 23,122 cells from experiment 1 and 26,611 cells from experiment 2 were retained. Data was merged into a single Seurat object for downstream analysis. To ensure that clustering would not be impacted by batch effects or biological variability we used the integration method implemented by Seurat v3. We integrated based on biological replicates using the SCT method to regress out cell cycle and mitochondrial associated genes prior to calculation of principal component analysis (PCA) and uniform manifold approximation projection (UMAP). Cell clusters were characterized by assessment of cell-specific genes and comparison with cell classification tool Single Cell Net.^22^ Cell prioritization was performed with the R package Augur (https://github.com/neurorestore/Augur) to identify populations that were most affected by PAH.^23^

Clusters corresponding to endothelial, stromal, immune, and epithelial cells were identified by *Cldn5, Col1a1, Ptprc, and Epcam*. These populations were re-clustered using the same normalization and integration approach to refine cluster identification and for subsequent analysis. Contaminating clusters were removed representing small populations with mixed canonical markers (eg. *Ptprc*+ cluster within the endothelial subset) likely representative of residual doublets not identified during previous steps.

### Differential gene expression and gene set enrichment analysis

Differential gene expression analysis was performed using the R package *muscat* (https://github.com/HelenaLC/muscat) to account for the available biological replicates,^24^ using standard workflow. Significant genes were identified with an adjusted p value <0.05 and a detection rate of at least 5% within the tested conditions. Gene set enrichment analysis was performed using R package *fgsea* (version 1.20.0) on the fold change ranked list of genes for a given condition tested. Gene sets were queried and annotated which comprised all GO terms, KEGG pathways, Reactome pathways, and the MSigDB Hallmark gene sets, which were acquired from the Molecular Signatures Database (v6).^25,26^ Significantly affected gene sets were filtered on an adjusted p-value of <0.05. The normalized enrichment score (NES) indicated the degree of up or down regulation of a given gene set. Volcano plots of DEGs were generated with r package EnhancedVolcano (version 1.12.0).

### Ingenuity pathway analysis

Pathway enrichment analysis was performed assessing only DEGs using Ingenuity Pathway Analysis (IPA) web-based software application (Qiagen). Standard workflows were followed within the core analysis function to identify gene sets of disease or biofunctions from the IPA library that were significantly enriched based on the input DEGs.^27^

### Transcription factor analysis

Transcription factor (TF) activity was predicted from available transcriptomic data using deCoupleR (version 2.0.1).^28^ We employed the weighted mean method using our gene expression data to infer TF activity and their targets from the DoRothEA curated network.^29^

### RNA velocity and trajectory inference analysis

We used scVelo^30^ as an input for CellRank^31^ to evaluate RNA velocity profiles of the combined vascular cells (ECs and stromal cells). We employed scVelo’s dynamical model to infer macrostates as an input for CellRank, which then performed RNA velocity and trajectory inferences.

### Immunofluorescent staining

OCT prepared lung sections were removed from −80°C freezer and thawed to room temperature and fixed in 4% paraformaldehyde solution for 20 minutes. Sections were washed three times in 1x PBS (15 minutes per wash) and then blocked with 5% goat serum in 0.5% Triton X-100/PBS (PBS-Tx) for one hour at room temperature. After blocking, samples were incubated in primary antibodies overnight at 4°C. The next morning samples were washed in PBS-Tx (3x, 15 minutes per wash) and incubated with secondary antibodies overnight at 4°C. The following morning samples were washed in PBS-Tx and prepared for confocal microscopy with mounting media containing DAPI. All images were captured using a laser scanning Zeiss 880 confocal microscope with Fast Airyscan (Zeiss) and processed with Aivia software. Primary antibodies with their respective concentrations were as follows Griffonia Simplicifolia Lectin I (GSL I) Isolectin B4 (1:25, Vector Laboratories, #FL-1201), mouse monoclonal anti-3G5 IgM (1:10, from Ke Yuan lab), FITC anti-αSmooth Muscle Actin mouse monoclonal antibody (1:300, Sigma Aldrich, #F3777), and rabbit anti-TM4SF1 antibody (1:25, Invitrogen, #PA5-21119).

### Human sample preparation

PAH patient and control lung samples were available from the Quebec Respiratory Health Network tissue bank (www.rsr-qc.ca). Paraffin sections (5µm thick) were deparaffinized and rehydrated as previously described using a series of xylene, ethanol, and water washes. Heat mediated antigen retrieval was performed by pressure cooking in citric acid-based antigen unmasking solution. Sections were washed in PBS with 0.1% Tween-20 (PBS-T) and blocked in 5% goat serum (Cedarlane #CL1200-500) in PBS-T for 1h at room temperature. Primary antibodies were incubated overnight at 4°C using TM4SF1 (ThermoFisher Scientific #PA5-21119, 1:200) and CD31 (Dako #M0823, 1:100) antibodies. The following day samples were washed 4x in PBS-T and incubated with secondary antibodies anti-Rabbit alexa fluor 488 (ThermoFisher Scientific #A11008, 1:500) and anti-mouse alexa fluor 594 (ThermoFisher Scientific #A11005, 1:500) for 1h at room temperature. Samples were washed in PBS-T, mounted using DAPI (4ʹ,6- diamidino-2-phenylindol) Fluoromount G mounting medium (Electron Microscopy Science) and imaged with an Axio Observer microscope (Zeiss).

### Statistical analysis

Data are presented as means +/- SEM. Non single cell statistical analyses were performed with GraphPad Prism v9 (GraphPad Software). Analysis of variance (ANOVA) was performed with multiple comparisons using Tukey correction used to identify statistically significant differences. P-values <0.05 were considered significant. For scRNA-seq experiments we used four animals per time point and statistical analysis was performed in line with recommendations for each analytical package used.

## Results

### Characterization of functional and structural vascular changes in the SU/CH PAH model

The SU/CH model exhibits many of the hemodynamic and pathological changes characteristic of human PAH, including complex arterial remodeling leading to the development of occlusive arterial lesions.^32^ To better define the stages in the development of PH, we evaluated animals at serial timepoints after the administration of SU and initiation of CH (**Figure 1A**). Significant elevation of RVSP was observed as early as 1-week (47mmHg; p = 0.056), plateauing at >100mmHg by 5-weeks (**Figure 1B**). Right ventricular hypertrophy (RVH) was increased by 3-weeks, persisting to 8 weeks (Fulton index >60%; **Figure 1C**). In small arterioles (<50µm), there was evidence of severe arteriolar remodeling (>50% lumen loss) by 5 and 8 weeks, compared to healthy lungs (**Figure 1D**). MicroCT analysis revealed a progressive loss of pulmonary arterial vasculature during PH progression, (**Figure 1E** **– H; Sup** **Figure 1**). The loss of the functional arterial bed primarily affected vessels <150µm in diameter, with evidence of significant arterial pruning as early as 1-week (**Figure 1F**), corresponding to the initial increase in arterial pressures, reaching a maximal loss of 65% at 5-weeks in vessels <200µm in diameter (**Figure 1G**). RVSP and lung arterial volume were negatively correlated (**Figure 1H**). Therefore, the rat SU/CH model is well suited to explore the cellular and molecular mechanisms during the development of PAH.

### Transcriptomic changes in global lung cell populations in PAH

Multiplexed single cell analysis was performed on lung samples from healthy controls, and 1-, 3-, 5-, 8-weeks post SU/CH initiation, incorporating four biological replicates per timepoint. Poor quality cells and those lacking a sample barcode or containing multiple barcodes were removed (**Sup** **Figure 2**). Additional doublet removal was performed with scDblFinder,^21^ and independent experiments (23122 cells Expt1 and 26611 cells Expt2) were merged, integrated, and aligned (**Sup** **Figure 3A, B**). Broad cell type annotations (endothelial, stromal, myeloid, lymphoid, and epithelial) were assigned to resulting clusters based on the expression of canonical markers and each were then subclustered for high resolution analysis (**Sup** **Figure 3C**). This resulted in the designation of 44 unique lung clusters (**Figure 2A**). Global UMAPs of all lung cell populations at each time point are presented in **Sup** **Figure 4**. At 1-week, transcriptomic changes were clearly evident in many cell populations, involving all lineages, but most evident in endothelial and myeloid cells (**Figure 2B, left)**. Surprisingly, at 5 weeks there was a remarkable and somewhat paradoxical ‘normalization’ of global transcriptional profiles, at a timepoint when the hemodynamic and vascular changes of PAH were reaching their peak (**Figure 2B, right**). Augur^23^ was used to identify cell populations most affected by SU/CH at 1-and 5-weeks relative to control cells. At 1 week, many cell clusters showed marked transcriptional changes, with the activated arterial (aA) ECs and a smooth muscle (SM)-like, *Acta2* positive pericyte populations demonstrating the most substantial changes from control conditions (**Figure 2C**). In contrast, at 5 weeks, there was an overall resolution of transcriptional changes with the notable exception of the aAECs (**Figure 2D**). Similarly, at early timepoints high numbers of differentially expressed genes (DEG) were observed across many different cell populations (**Figure 2E**), with the greatest increases seen in immune cells, particularly transitional monocytes (>400 DEGs), and various endothelial populations at 1 week. However, again there was a progressive decrease in the number of DEGs over time, with the exception of aAECs, which continued exhibit ∼150 DEGs for up to 8 weeks. Expansion in relative cell numbers was seen in several populations including the arterial and aAEC clusters, and lymphatic ECs clusters, as well as some monocyte (alveolar, interstitial, and transitional macrophages), neutrophil and NKT cell clusters (**Sup Figures 5-8**). However, SM-like *Acta2*^+^ pericytes exhibited the most robust relative increase in cell number beginning at 3 weeks post SU/CH coinciding with the advent of arterial remodeling (**Sup Figure 5**). For the remainder of this study, we focus mainly on the endothelial and stromal populations, which showed the most marked overall changes in the global analyses. However, relevant details of cell annotation and transcriptomic changes for myeloid, lymphoid, and epithelial cells are provided in **Sup Figures 9-11**.

**Figure 2:**
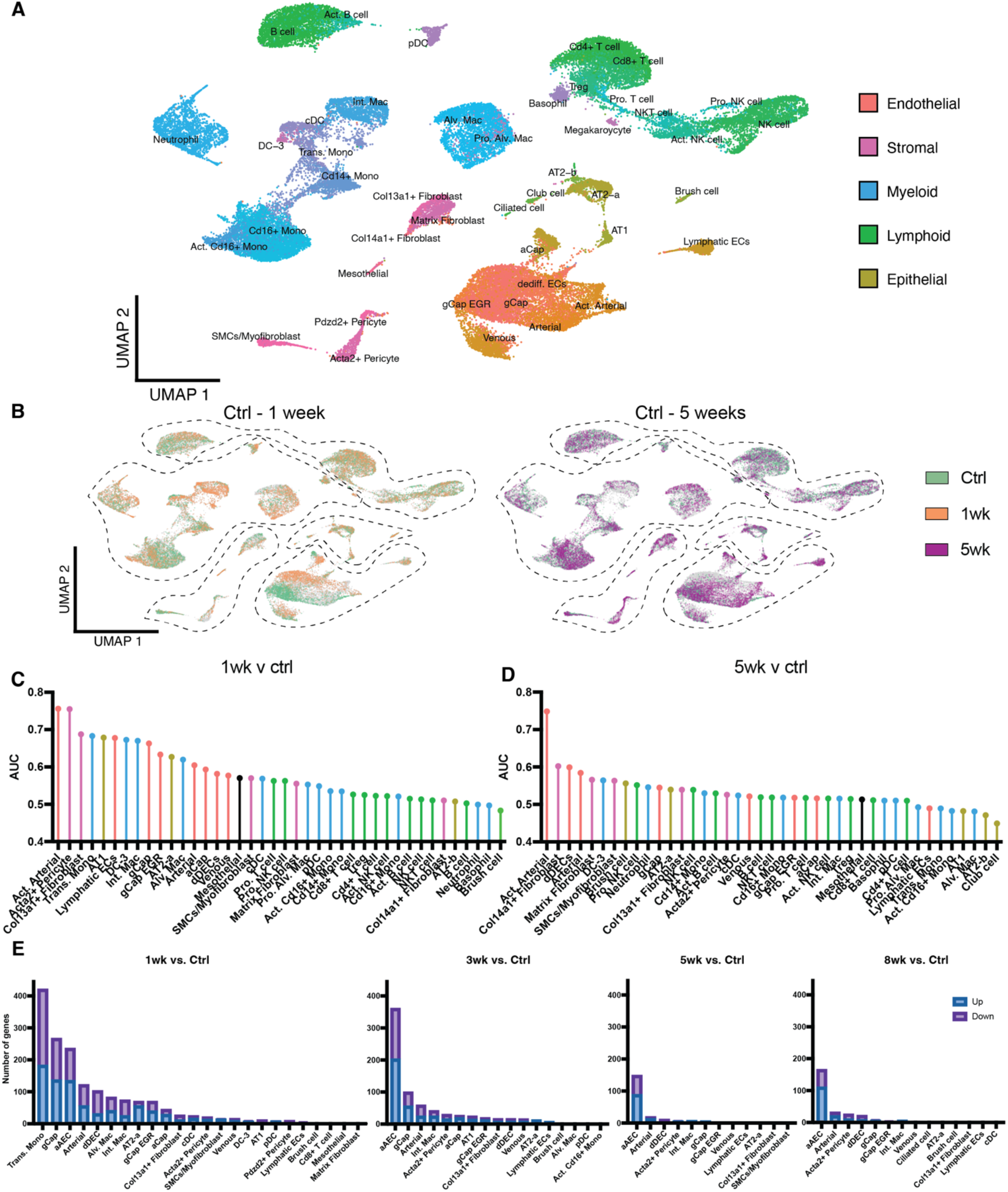
Multiplexed single cell transcriptomic atlas of lung cells during PAH progression. A) 44 unique lung clusters were identified and integrated into the global uniform manifold approximation and projection (UMAP). B) Global UMAPs colored by timepoint with control (green) compared with 1-week (orange) and 5-weeks (purple). Cell prioritization was performed using a machine learning algorithm to identify clusters that were most affected between 1-week and control (C) and 5-weeks and control (D). At 1-week major transcriptomic changes indicated by an increase in area under the curve (AUC) were seen in many populations with the greatest increases in activated arterial ECs (aAECs) and *Acta2*+ pericytes. However, at 5-weeks there was a general reduction in transcriptomic changes, with the exception of the aAEC population E) Number of all non-zero differentially expressed genes (DEGs) in lung cell populations at 1-, 3-, 5-, and 8-weeks post SU compared with control (healthy). High numbers of DEGs were seen in transitional macrophages and many EC clusters at 1-week, but only aAECs displayed a persistently high numbers of DEGs at later timepoints. Single cell data was obtained from n = 4 animals per timepoint, multiplexed using unique barcodes, pooled and subjected to library construction using 10x-Genomics.

**Figure 3:**
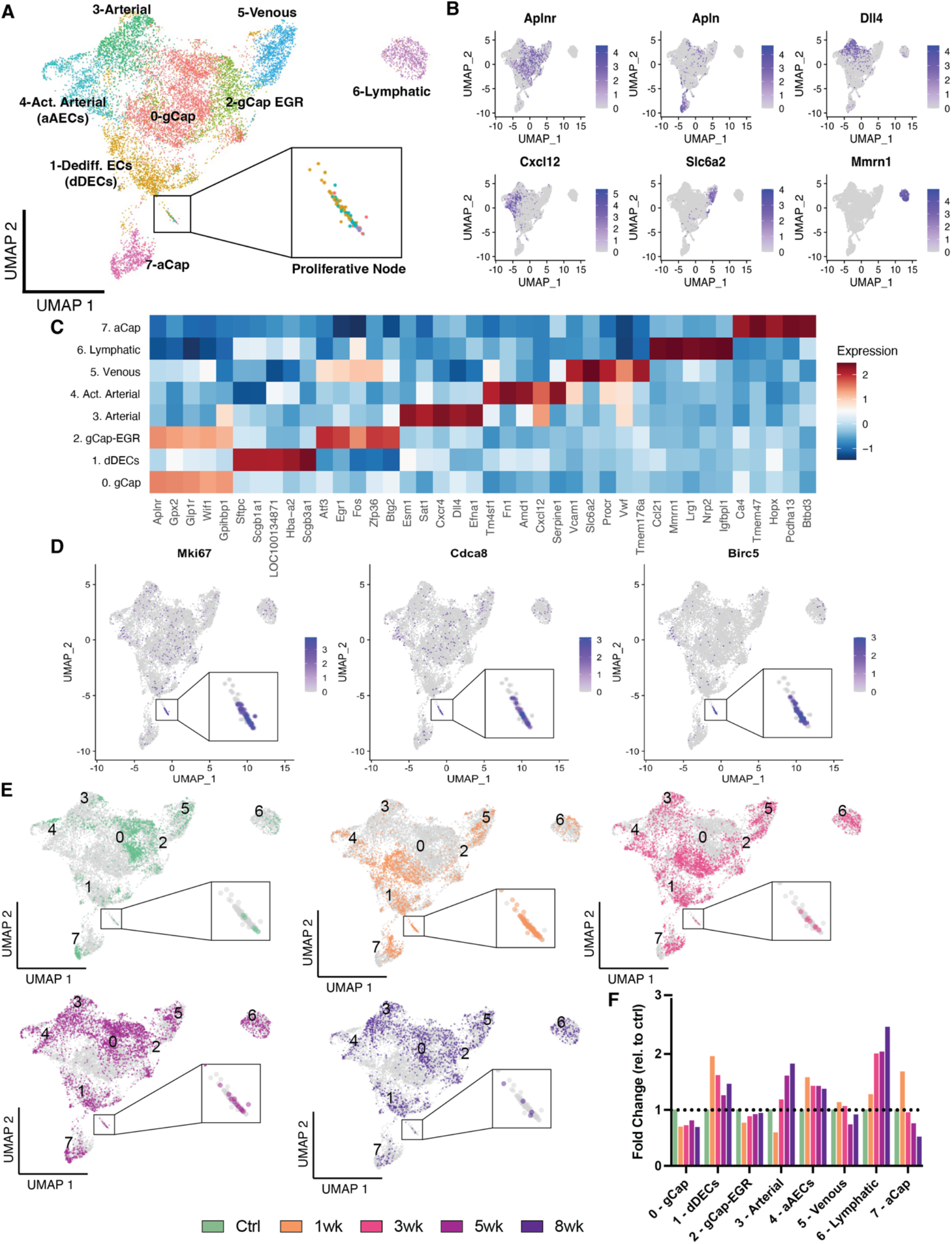
Changes in endothelial cell populations during PAH progression. A) Uniform manifold approximation and projection (UMAP) representation of all endothelial populations at all sampled timepoints showing 8 distinct cell types and a proliferative cell node enlarged in the box. B) Feature plots highlighting genes typically associated with distinct endothelial cell (EC) populations including: *Aplnr* (gCap), *Apln* (aCap), *Dll4* (arterial), *Cxcl12* (‘activated’ arterial [aAECs] and arterial), *Slc6a2* (venous) and *Mmrn1* (lymphatic). C) Heatmap showing the top 5 differentially expressed genes distinguishing each EC population. D) Feature plots showing the distribution of proliferation-related genes, *MKi67*, *Cdca8* and *Birc5*, largely within the proliferative node, highlighted and enlarged in the box. E) UMAP of cells present at each timepoint demonstrating transcriptomic shifts compared to control at 1-, 3-, 5-and 8-weeks. F) Fold change of each EC populations relative to their control levels. Early and persistent increases were seen in the relative size aAECs and ‘dedifferentiated’ (dD) ECs, whereas as arterial and lymphatic ECs showed a later expansion.

**Figure 4:**
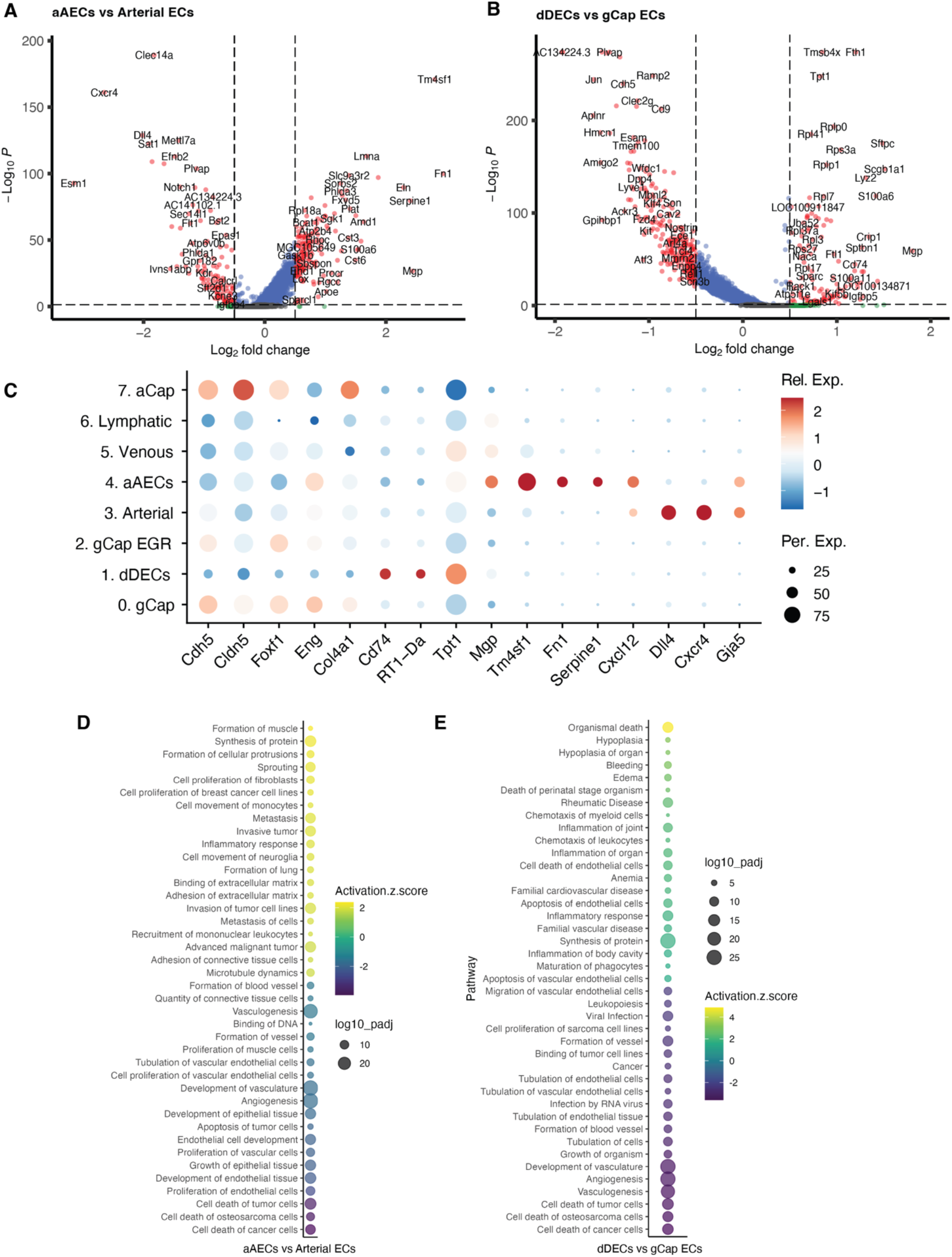
Transcriptomic signature of disease-specific activated arterial and dedifferentiated EC populations. Volcano plots for all differential expressed genes between A) aAECs and arterial ECs and B) dDECs and gCap ECs including all timepoints. C) Dot plot showing relative expression of genes of interest between all EC clusters. dDECs exhibited the lowest expression of *Cldn5* and *Cdh5*, and the highest expression of *Cd74*, *RT1Da* and *Tpt1*, whereas aAECs showed the highest expression of *Tm4sf1*, among other genes associated with an activated phenotype and low expression of typical arterial genes (*Dll4* and *Cxcl4*). D) Ingenunity pathway analysis (IPA) comparing aAECs and arterial ECs showing the top 15 significantly up and down regulated gene sets. aAECs exhibited upregulation of gene sets associated with cancer cell proliferation, migration and invasion, with downregulation of gene sets for angiogenesis, vascular development and apoptosis. E) IPA comparing dD and gCap ECs showing the top 15 significantly up and down regulated gene sets, demonstrating upregulation of gene sets in dDECs associated with inflammation and apoptosis and downregulation of gene sets associated with vascular development and angiogenesis.

**Figure 5:**
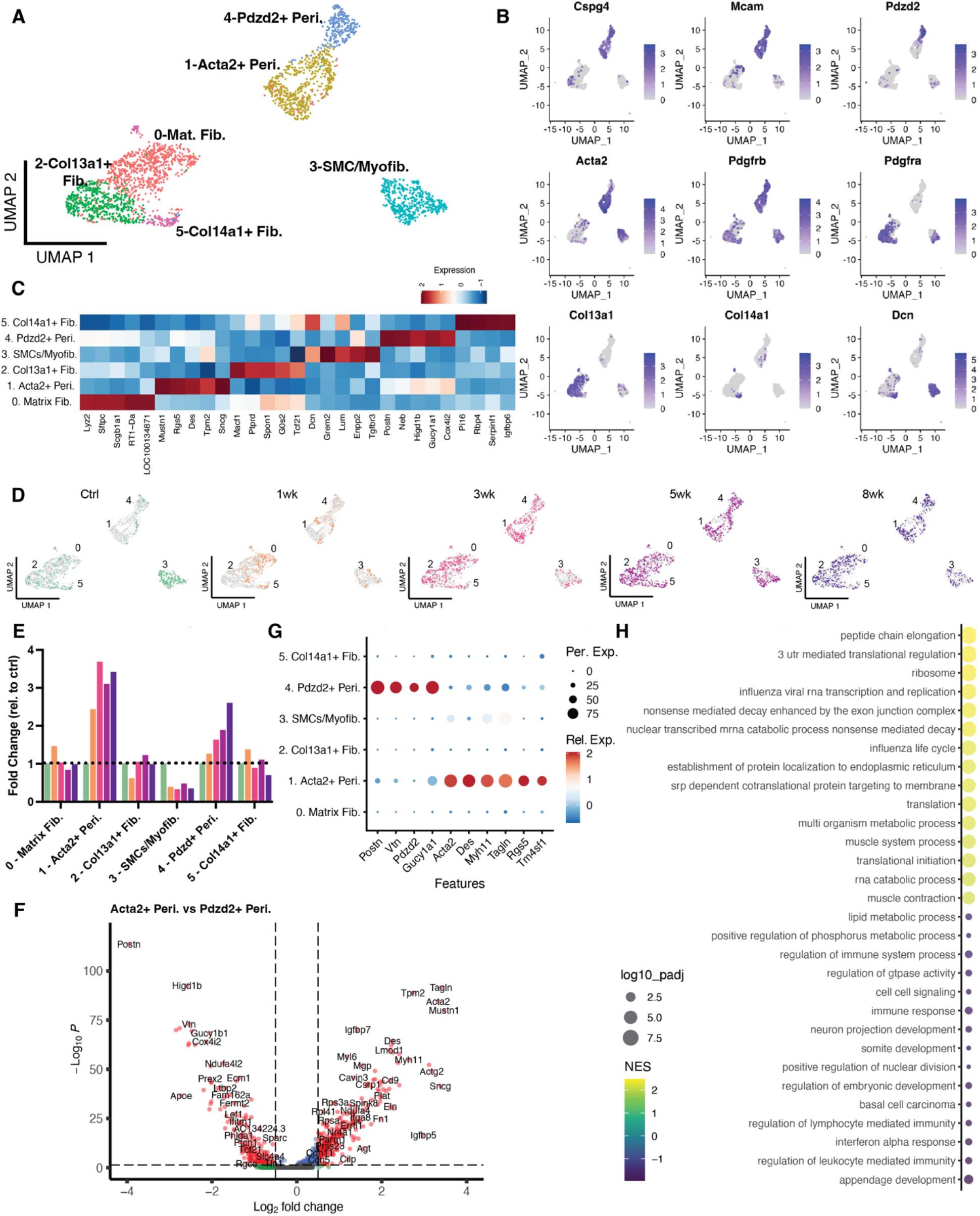
Changes in stromal cell populations during PAH progression. A) Uniform manifold approximation and projection (UMAP) representation of all stromal populations at all sampled timepoints showing 6 distinct cell types. B) Feature plots highlighting genes typical of distinct stromal populations including: pericytes (*Cspg4, Mcam, Pdgfrb*); fibroblasts (*Pdgfra, Col13a1, Col14a1*); smooth muscle cells (*Acta2*); and myofibroblasts (*Dcn*). C) Heatmap of top 5 differentially expressed genes distinguishing each stromal population. D) UMAP portraying the presence of cells from each stromal population at each timepoint, showing a marked expansion of the pericyte populations by 3-weeks. E) Fold change of each stromal population relative to their respective controls demonstrating a marked relative increase in the ‘smooth muscle’ (SM)-like *Acta2*+ pericytes and less so the ‘classical’ pericyte population during PAH progression. F) Volcano plot of all differentially expressed genes (DEGs) between SM-like pericytes and classical pericytes. G) Dot plot showing relative expression of genes of interest between different pericyte populations. H) Gene set enrichment analysis with the top 15 up and down regulated gene sets between SM-like pericytes and classical pericytes.

### Emergence of disease-specific endothelial populations during PAH progression

Sub-clustering of endothelial cells produced 8 distinct cell clusters encompassing all expected populations including arterial, venous, lymphatic, general capillary (gCap) and aerocytes (aCap) ECs (**Figure 3A**), as identified based on expression profile of commonly used markers ^7,33,34^ such as, *Dll4*, *Slc6a2*, *Mmrn1*, *Aplnr*, *Apln*, among many others (**Figure 3B and C**). Interestingly, arterial ECs consisted of two distinct populations, a ‘classical’ arterial cluster and the ‘activated’ aAEC cluster, which is further characterized below. Similarly, the gCap EC population was made up of two clusters; a typical gCap EC population and another cluster characterized by expression of early response genes (ERG), such as *Fos* and *Egr*, as was previously described.^19^ Of interest, the gCap-EGR cluster shared expression of some of these early response genes with the venous ECs (**Figure 3C**), suggesting possible zonation changes across the capillary-venous axis.^33^ Finally, a novel EC population (cluster 1) was identified, termed ‘de-differentiated’ ECs (dDECs) that exhibited an atypical endothelial gene expression profile with some of their top distinguishing genes related to nonendothelial lineages. Again, this cluster is further characterized below.

Using proliferative markers, including *Mki67*, *Cdca8*, and *Birc5*, evidence of proliferation was seen in cells mainly located in a distinct ‘proliferative’ node (**Figure 3D**), representing cells from several clusters, in particular dDECs and aAECs (**Figure 3A**). Endothelial UMAPs for each timepoint show the major transcriptomic changes between healthy (control) and early stages (1 and 3 weeks), or the later stages (5 and 8 weeks) of PAH progression (**Figure 3E**). Phenotypic changes were most evident in the gCap at weeks 1 and 3 but returned to the baseline distribution at weeks 5 and 8. This was accompanied by the marked expansion in clusters 1 and 4 (dDECs and aAECs, respectfully) at 1 week which persisted to week 8. The fold-change relative to control in each EC cluster, proportional to the total EC population, is shown in **Figure 3F**. There was an early and sustained increase in the dDECs and aAEC populations, with later expansion in arterial and lymphatic ECs. Again, this supports an important role for the aAECs and dDECs as disease-specific endothelial populations during the onset and progression of PAH.

### Differential gene expression analysis in activated arterial and dedifferentiated EC populations

To better understand the distinct transcriptomic profiles within these populations, we compared changes in gene expression (including all timepoints) between arterial ECs and aAECs and between gCap and dDECs since these populations were closely related on the UMAP (**Figure 4A and B**, respectively). aAECs were characterized by reduced expression of typical arterial genes,^7^ including the notch ligands (*Dll4* and *Notch1*), gap junction proteins (i.e., *Gja5*), signaling molecules (*Efnb2* and *Efna1*), and chemokine receptors (i.e., *Cxcr4*) (**Figure 4A**). Moreover, they showed increased expression of the transitional extracellular matrix (ECM) proteins, such as fibronectin (*Fn1*), Matrix gla protein (*Mgp*) and elastin (*Eln*), and signaling molecule stromal derived factor 1 (*Cxcl12*), all consistent with an activated endothelial phenotype. Of particular interest, transmembrane 4L6 family 1 (*Tm4sf1*), has been strongly implicated in cancer cell growth,^35^ The differences in gene expression between gCap and dDECs were consistent with a de-differentiated state of dDECs with decreases in endothelial tight junction genes, VE-cadherin (*Cdh5*) and claudin 5 (*Cldn5*), as well as typical gCap markers such as plasmalemma vesicle associated protein (*Plvap*) and apelin receptor (*Aplnr*) (**Figure 4B**). Interestingly, numerous genes associated with ribosomal proteins (*Rpl*) were upregulated in dDECs, which is a feature of a hyper-transcriptomic state characteristic of progenitor cells,^36^ together with genes associated with antigen presentation and inflammation (*Cd74*, *RT1-Da*), and tumor protein, translationally-controlled 1 (*Tpt1*) (**Figure 4B**), an oncogene and proinflammatory factor previously implicated in PAH.^37,38^ The relative expression in selected genes between all EC populations are depicted in **Figure 4C**, showing the relative overexpression of genes associated with an activated endothelial state in aAECs, in particular *Tm4sf1*, whereas dDECs exhibited a predominance of genes associated with antigen presentation, while displaying lower expression of typical EC tight junction genes. By ingenuity pathway analysis (IPA), aAECs were enriched for biofunctions associated with cell proliferation, protein synthesis, invasion and inflammation consistent relative to classic arterial, consistent with a ‘cancer-like’ phenotype, together with reduced expression of gene sets associated with angiogenesis and vascular development (**Figure 4D**). In contrast, dDECs showed enrichment of gene sets for inflammation, protein synthesis, cardiovascular disease and cell death/apoptosis compared to gCap ECs (**Figure 4D and E**), consistent with a role vascular disease.

### Changes in stromal cells in response to PAH

Next, we re-clustered all stromal cell populations (fibroblasts, pericytes, smooth muscle cells) into 6 unique populations, including matrix fibroblasts, *Col13a1*+ fibroblasts, *Col14a1*+ fibroblasts, *Acta2+* pericytes, *Pdzd2*+ pericytes, and smooth muscle cells (SMCs)/myofibroblasts (**Figure 5A**). Fibroblasts were identified based on *Pdgfra* and *Tcf21* expression, and differential expression of *Col13a1* and *Col14a1*; while pericytes were identified by expression of *Pdgrb*, *Csgp4,* and *Mcam* (**Figure 5B and C**).^9,39^ Pericytes were further distinguished into ‘classical’ pericytes uniquely expressing *Pdzd2*, *Postn*, *Gucy1b1* and ‘SM-like’ pericytes based on expression of contractile markers (*Acta2, Des*) (**Figure 5B and C**), as previously described by Hurskainen et al.^9^ SMCs and myofibroblasts were observed in a mixed population expressing *Acta2* and *Dcn*. Interestingly, both pericyte populations showed marked expansion during PAH progression, consistent with a possible contribution to arteriolar remodeling, in PAH whereas there was little change in the global proportion of other stromal cell populations (**Figure 5D and E**).

We then compared transcriptomic profiles of ‘classical’ and SM-like pericytes (at all timepoints). The SM-like pericytes exhibited high expression of genes associated contractile function including, *Acta2*, *Des*, *Myh11*, and *Mustn1*, as well as reduced expression of some typical pericyte genes, including *Postn, Gucy1a1, Pdzd2* and *Vtn* compared to classical pericytes (**Figure 5F and G**). Like aAECs, SM-like pericytes were somewhat enriched in *Tm4sf1* (**Figure 5G**), consistent with a proliferative and invasive phenotype. Gene set enrichment analysis (GSEA) comparing SM-like pericytes to classical pericytes showed that SM-like pericytes exhibited robust enrichment of gene sets associated with contraction, muscle differentiation and translation, while gene sets associated with GTPase activity, lipid processing and organ development were decreased (**Figure 5H**). Therefore, these data are consistent with the SM-like pericytes possibly contributing to arteriolar remodeling and distal muscularization in PAH.

### Temporal profile of endothelial transcriptomic changes in aAECs during PAH progression

By performing scRNA-seq at multiple timepoints in the SU/CH model, we were able to discern changes in gene expression profiles during the development of the PAH phenotype. Volcano plots prtray DEGs within the aAEC cluster between control animals and SU/CH rats at weeks 1 or 5 (**Figure 6A and B**, respectively). A heatmap showing the top 20 up or down regulated genes for the aAEC cluster at each timepoint revealed three distinct profiles (**Figure 6C**). The first was represented by an early increase in gene expression peaking at 1 week, with a partial normalization by 5 to 8 weeks. These were largely cell-cycle associated genes, including *Birc5*, *Cdc20*, *Cdkn2c*, *Cdkn3*, and *Cdk1* (**Figure 6C**). Interestingly, a similar pattern of gene expression was observed in the dDECs at 1 week, consistent with the increase in EC proliferation observed in both populations (**Figure 3D, E**). A second pattern was unique to aAECs and was represented by genes that showed a sustained increase in expression compared to control at all timepoints and this included a number of genes which have previously been implicated in PAH such as, *Ramp1*,^40^ *Trpv4*,^41^ *Jag1*,^42^ and *Pdgfa*^43^. Finally, a third pattern was made up of genes strongly expressed at baseline by both classical arterial ECs and aAECs, largely involved in angiogenesis and vascular development, such as *Dll4*, *Notch1*, *Efna1*, and *Erg1*, which were selectively downregulated in aAECs during PH development (**Figure 6C**). Not unexpectedly, *Cyp1a1* and *Cyp1b1*, which mediate drug metabolism,^44^ were upregulated in all EC populations after treatment with SU, with the exception of lymphatic ECs (**Figure 6C**). Ingenuity pathway analysis (IPA) of changes in gene expression in aAECs between control and PAH timepoints confirmed the upregulation of gene sets associated with increased cell proliferation and cancer-like cell growth in aAECs, with downregulation of gene sets associated with angiogenesis, vasculogenesis and cell death (**Figure 6D**). Together, these results point to aAECs as a unique lung EC population exhibiting a transcriptional signature consistent with a growth dysregulated state, associated with decreased reparative and angiogenic capacity, and likely playing an important role in arterial remodeling in PAH.

**Figure 6:**
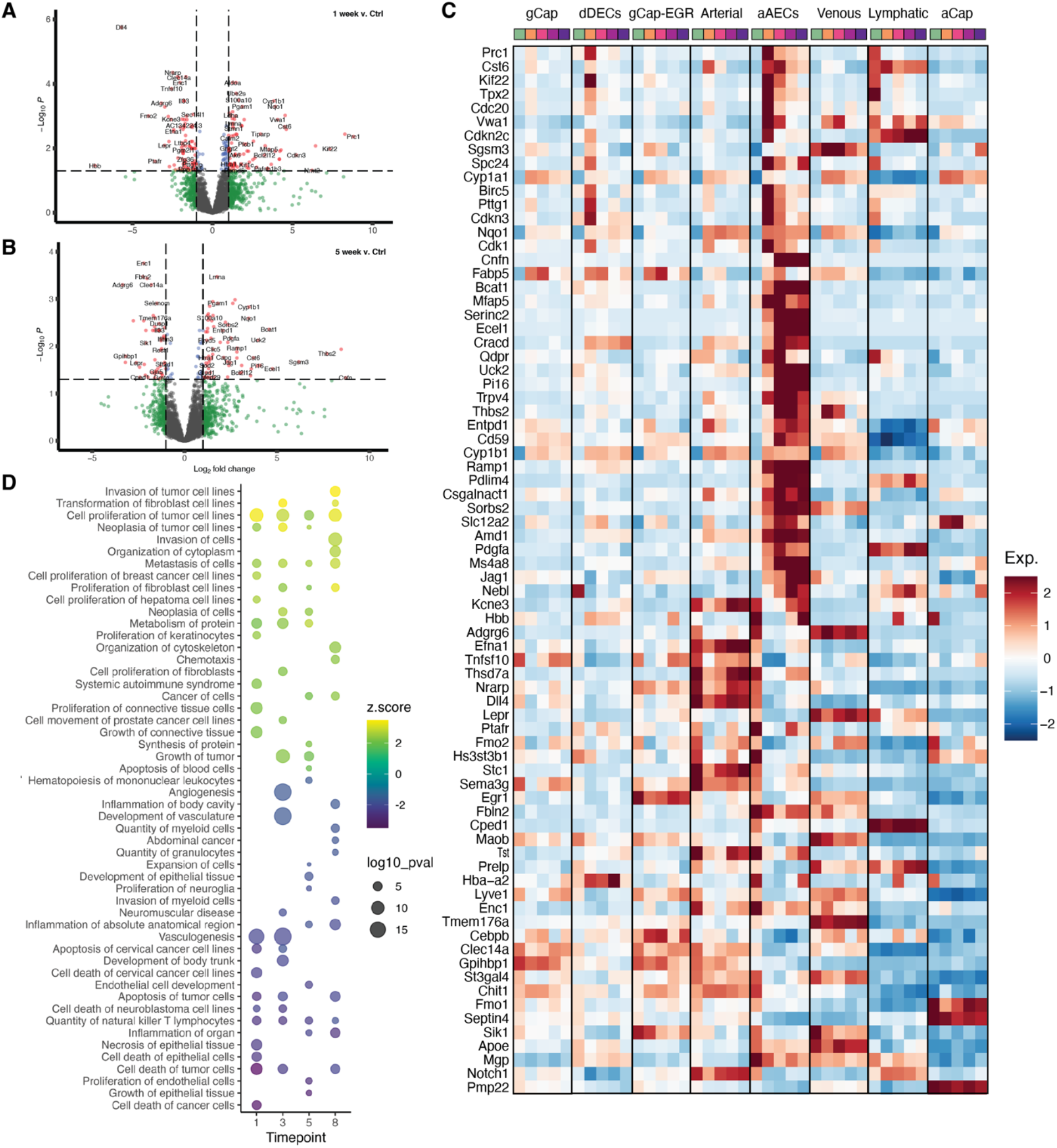
Endothelial transcriptomic changes during onset and progression of PAH. Volcano plots showing all DEGs for aAECs vs control at 1-week (A) and 5-weeks (B). C) Heatmap showing the top 20 up and downregulated DEGs for aAECs at each timepoint relative to control. Three transcriptional patterns can be discerned: i) peak increase at 1-week with partial normalization at later timepoints, ii) persistent elevation at all timepoints and iii) transient or sustained reduction in gene expression. D) Ingenuity pathway analysis (IPA) for aAECs comparing each timepoint to the control, showing the top up or downregulated disease or biofunction gene sets for each timepoint.

### Spatial localization of *Tm4sf1*-expressing activated arterial ECs

Both the aAECs and SM-like (*Acta2*^+^) pericytes exhibited expression of *Tm4sf1*, raising the possibility that this surface protein may be used as a marker of these populations. Therefore, we examined *Tm4sf1* expression across all 44 lung cell populations, and found that *Tm4sf1* was largely restricted to arterial and venous ECs, as well as pericytes and mesothelial cells, with by far the greatest expression in aAECs (**Figure 7A**). We then used immunofluorescent staining for TM4SF1, and co-staining using isolectin B4 (ECs), 3G5 (pericytes) or smooth muscle actin (SMA; smooth muscle cells), to identify the anatomic location of these cell populations in the healthy and diseased lungs. In control lung samples, TM4SF1 expression was localized mainly to 3G5^+^ pericytes surrounding alveolar capillaries, with sparse staining of ECs (**Figure 7B**). However, at 5 weeks post SU/CH, there was abundant TM4SF1 staining largely associated with ECs, identified by isolectin B4, localized mainly to severely remodeled arterioles with accumulation of TM4SF1^+^ ECs seen within the vascular lumen, consistent with a direct role for the aAECs in occlusive arteriopathy (**Figure 7C**). Interestingly, in the diseased lung, 3G5 staining was tightly associated with SMA^+^ bands of neomuscularization, suggesting a role for the SM-like pericytes in distal arteriolar medial remodeling. Notably, TM4SF1 staining was reduced in these SMA positive regions, possible indicating that as SM-like pericytes differentiate further towards a mature smooth muscle phenotype, they may lose the expression of this ‘progenitor’ cell marker. We also assessed the spatial distribution of TM4SF1 expression in human lung samples from control and PAH patients (**Figure 7D**). In the control samples, TM4SF1 staining could be seen surrounding small arterioles, but was not associated with ECs (CD31 staining). In contrast, in PAH samples there was abundant TM4SF1 staining, localized to the perivascular region but also adjacent to the intima, and clearly associated with ECs, with a relative paucity of staining in the intervening SMC layer (**Figure 7D**). Together these results confirm that TM4SF1 positive ECs are associated with complex arterial remodeling in PAH, together with SMA^+^ pericytes, consistent with a pathological role of aAECs and SM-like pericytes in this disease.

**Figure 7:**
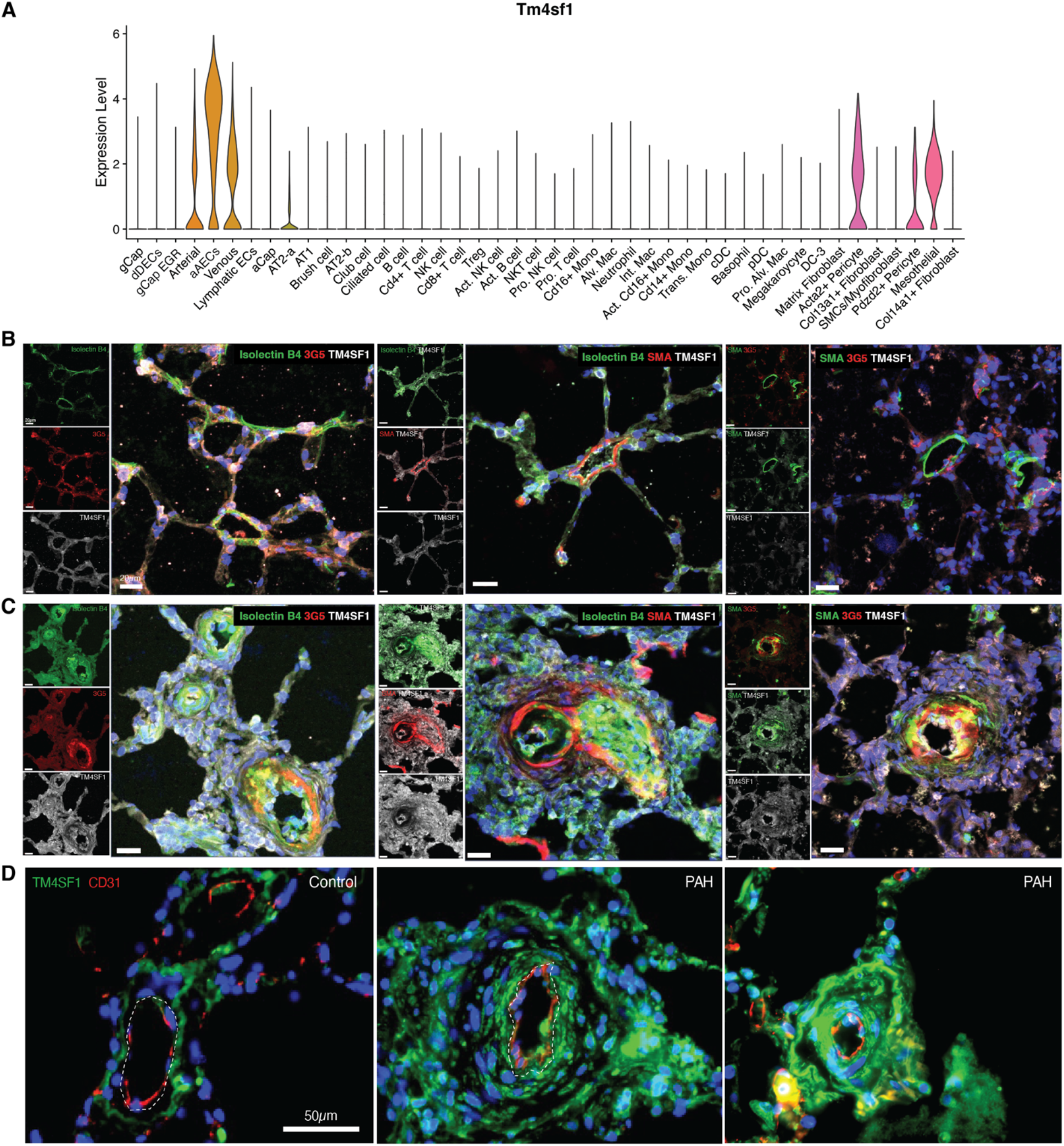
Tm4sf1 as a marker for aAECs and SM-like pericytes in arterial remodeling. A) Violin plots showing *Tm4sf1* the highest expression in aAECs, with lower levels in arterial and venous ECs, SM-like pericytes and mesothelial cells. Immunofluorescent staining for TM4SF1 (white), isolectin B4 (ECs, green), 3G5 (pericytes, red) or SMA (smooth muscle cells, red) in healthy lungs (B) or at 5-weeks in the SU/CH severe PAH model (C). D) Immunofluorescent staining of lung sections from control subjects and PAH patients for TM4SF1 (green), co-stained for ECs (CD31, red). Scale bar represents 20μm (B and C) and 50μm (D).

### RNA velocity of endothelial and stromal cells during SU/CH progression

Endothelial to mesenchymal transition (EndMT) has been previously implicated in PAH.^45^ To explore a potential role of cell fate transitions in the development of PAH we employed RNA velocity analysis, using scVelo dynamical modelling,^30^ specifically addressing population dynamics within the EC and stromal clusters based on their importance in arterial remodeling. Notably, RNA velocity analysis demonstrated vectors originating in the arterial and aAEC populations passing through the dDECs cluster and directed to fibroblast populations (**Figure 8A**), consistent with the possibility of EndMT contributing to vascular stromal cells in this PAH model. Velocity vectors were also observed from the classical (*Pdzd2*^+^) pericytes towards the SM-like (*Acta2*^+^) pericytes, consistent with pericyte differentiation towards a smooth-muscle cell fate as a mechanism of distal muscularization. There were also vectors from the gCap-EGR cluster leading to the venous ECs, again supporting the possibility of zonation of this cluster towards the venous side of the alveolar capillary. The possibility of EndMT was further supported by a latent time analysis which provides an estimate of a cell’s internal clock, from progenitor fates (approaching 0) to terminally differentiated cells (approaching 1). This model identified aCap and gCap ECs, *Col13a1*^+^ fibroblasts, and classical (*Pdzd2*^+^) pericytes as being close to ‘terminal’ states, while aAECs, dDECs, and SM-like pericytes were closer to a progenitor state (**Figure 8A’**). We then inferred transcription factor (TF) activities from gene expression data for the EC clusters using deCoupleR.^28^ The location of dDECs (EC cluster 1) in the endothelial UMAP is shown in **Figure 8B**. TFs classically associated with EMT^46^ including *Zeb1*, *Zeb2*, *Snai2* had predicted activity in dDEC cluster, as well as some other EC populations, whereas *Twist1* was seen mainly in gCap-EGR ECs (**Figure 8C**). The top 40 most variable TFs are presented in **Figure 8D**. Importantly, dDECs exhibited very low expression of the endothelial-restricted transcription factor, ETS-related gene (*Erg*), a principal determinant of EC identity,^47^ as well as the Friend Leukemia Integration 1 Transcription Factor (*Fli1*), another member of the ETs family, which are both downregulated in EndMT.^48^ As well, dDECs exhibited elevated activity associated with *Nr2f2*^49^ and *Zeb2,*^46^ both which have been implicated in EndMT, further supporting a potential role for this relatively de-differentiated EC population in EndMT during PAH progression (**Figure 8D, insert**). Interestingly, gCap-EGR and venous ECs shared a number of TFs (**Figure 8D**), further supporting zonation between these populations based on similarities in gene expression as shown in **Figure 3C**.

**Figure 8:**
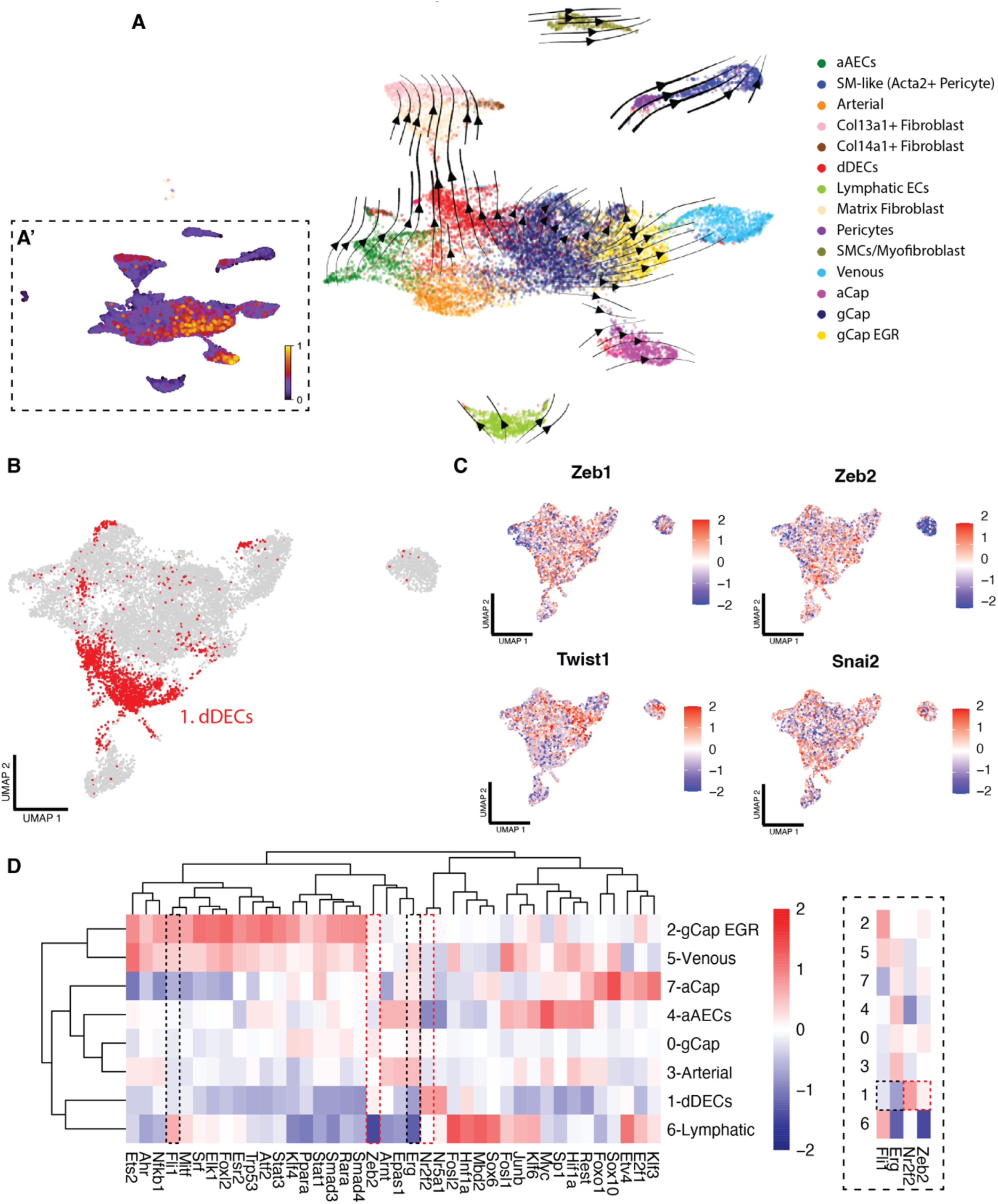
RNA velocity analysis indicates endothelial to mesenchymal transition. A) RNA velocity analysis of endothelial and stromal cell populations using scVelo’s dynamical model and CellRank demonstrate velocity vectors from aAEC and arterial EC populations, going through the dDEC cluster to fibroblast populations. RNA velocity vectors are also seen between classical pericytes SM-like (*Acta2*+) pericytes. Inset (A’) highlights the latent time analysis which estimates cellular differentiation and shows that gCap, aCap, fibroblasts, and classical pericytes approximate a more terminal differentiated state, while aAECs, dDECs and SM-like pericytes exhibit a more progenitor-like state. B) EC subset UMAP showing the distribution of dDECs in red. C) UMAP showing the distribution of typical epithelial to mesenchymal transition (EMT) related transcription factor (TF) activities in the EC populations. E) Heatmap of the top 40 most variable TFs in the different EC populations. The insert highlights reduced activity of *Erg* and *Fli1*, ETS family TFs that control endothelial identity, within cluster 1 (dDECs) and increased activity of *Nr2f2* and *Zeb2*, which have been implicated in EndMT.

## Discussion

PAH is characterized by marked pruning of the pulmonary arteriolar bed which, together with complex arterial remodeling, leads to progressive increases in pulmonary vascular resistance; yet the mechanisms underlying these dramatic vascular changes remains unclear. The rat SU/CH model of PAH provides an ideal opportunity to study functional, structural, and molecular changes during the onset and development of PAH, rather than only at late stages of disease when typically, clinical lung tissue is available.

Using multiplexed scRNA-seq we analyzed four timepoints, spanning early to late disease, comparing SU/CH to healthy control animals. Major transcriptomic changes were observed in nearly all lung cell populations at 1-week post SU/CH, including immune, stromal, and vascular cells. Notably, ‘transitional’ monocytes, exhibiting characteristics between those of classical and interstitial monocytes, were among the top five most transcriptionally affected in the PAH model based on the Augur machine learning algorithm and exhibited the greatest number of DEGs at week 1. This is consistent with an important early inflammatory response to injury induced by VEGFR2 inhibition coupled with hypoxia. However, at later timepoints, there was substantial normalization in gene expression profiles in nearly all lung cell populations, with the notable exception of the ‘activated’ arterial EC (aAECs) cluster. This finding was surprising considering that the major pathological and hemodynamic changes in this model manifest at later time points. In particular, the lack of persistence of a transcriptional signature for inflammation is remarkable given the abundant evidence that implicates immune mechanisms in the pathogenesis of PAH.^50^ However, this may in part reflect overrepresentation of cells from the distal parenchyma and microcirculation in the lung isolate, which account for by far the greatest number of cells after dispersion, whereas larger vessels, which exhibit more marked pathologically inflammatory changes, make up only a small proportion of total cells. Nonetheless, we did observe some intriguing immune signals including a persistent, albeit modest, increase in DEGs in interstitial macrophages and a robust expansion of the NKT population in later stages of disease. This is consistent with a previous single cell transcriptomic analysis of large pulmonary arteries from patients with severe PAH, which showed a significant increase in NKT cells, along with T cells and neutrophils, compared to control.^14^

In contrast, persistent and robust changes in transcriptional activity were seen selectively in one endothelial cluster, aAECs, suggesting that this population may be playing a central role in the progression of vascular disease in this model. Since the effects of SU and hypoxia would no longer be operative at later timepoints beyond 3 weeks, this raises the question of what drives the persistent transcriptomic response in the aAEC cluster. It has been reported that complex arterial remodeling in this model is dependent on perturbed pulmonary hemodynamics,^6^ likely due to pathological levels of shear stress, which would have the greatest impact on the distal arteriolar bed.^51^ Using micro-CT, we demonstrated a progressive loss of arteriolar volume in the SU/CH model, which was significant as early as 1 week, preceding any evidence of occlusive arterial remodeling. This suggests that early arteriolar loss, probably as a direct consequence of widespread EC apoptosis, may be the primary mechanism for the initial hemodynamic abnormalities in this model. As well, since in a dispersed lung cell preparation, distal arteriolar ECs will greatly outnumber those from larger arteries, the ‘arterial’ EC populations that we identified by scRNA-seq would be predominately arteriolar in origin. Thus, by virtue of being largely derived from the distal arteriolar bed, aAECs would have been among the cells the most affected by the abnormal shear forces associated with developing PH,^51^ which provides a likely stimulus for the persistent transcriptomic activity seen in this population. Indeed, this EC population may be the link between perturbed pulmonary hemodynamics and complex arterial remodeling that was previously described.^6,52^

At the 1 week timepoint, aAECs and dDECs exhibited a similar transcriptional signature characterized by the transient expression of genes related to cell proliferation and repair. However, at later time points (5 and 8 weeks), only aAECs exhibited a persistent dysfunctional gene expression profile that was consistent with dysregulated cell growth and reduced capacity for vascular repair and angiogenesis. Thus, using single cell transcriptomic analysis we have been able to map the origin of hyperproliferative and apoptosis-resistant ECs which have long been implicated in occlusive arterial remodeling in PAH.^53^ We have also demonstrated that aAECs can be distinguished from other EC populations by high expression of *Tm4sf1*, a trans-membrane protein which is also found in many cancer cells.^35^ Using immunofluorescence staining, we were able to show that TM4SF1^+^ ECs were abundant in complex arterial lesions in the SU/CH model and as well as in lung sections from patients with PAH, particularly in regions of occlusive arteriopathy, consistent with a dominant role for this EC population arterial remodeling. Interestingly, this marker was also expressed by the SM-like pericytes and, in the healthy lung, TM4SF1 staining was associated with pericytes surrounding alveolar capillaries. However, in the diseased lung, 3G5^+^ muscularized regions of remodeled arteries exhibited lower TM4SF1 staining, consistent with their differentiation to a more mature SMC phenotype. This finding also suggests that the SM-like pericyte cluster, which by RNA velocity analysis was derived from the classical Pdzd2^+^ ‘classical’ pericytes, may contribute importantly to distal arterial muscularization in PAH, as has been previously suggested.^54–56^

Hong et al. identified a similar arterial EC cluster expressing *Tm4sf1* in both the rat monocrotaline (MCT) model and SU/CH model in late-stage PH.^11^ However, the published transcriptomic profile of this *Tm4sf1*^+^ EC population included expression of genes characteristic of capillary (*Ednrb*), and venous (*Slc6a2, Icam1*), in addition to arterial ECs (*Gja5*). Indeed, we also found that *Tm4sf1* was expressed by venous, but not capillary ECs, but at much lower levels than in aAECs, which also uniquely demonstrated a persistent transcriptomic signature of dysregulated cell growth. As well, we also showed that ECs within occlusive arterial lesions in both the SU/CH model and human PAH patients strongly expressed TM4SF1, consistent with a key role for this population in complex arterial remodeling. While TM4SF1 may be a useful marker to identify aAECs, it is also possible that it may contribute to growth dysregulation.

TM4SF1 is a trans-membrane protein belonging to the tetraspanin superfamily that regulates numerous signaling pathways modulating cell development, activation, growth, and motility.^35^ Increased *Tm4sf1* expression has been observed in cancer cells, particularly in breast, lung and ovarian cancers,^35^ and is a promising target for novel cancer therapies.^57–59^ Therefore, similar strategies to antagonize or silence *Tm4sf1* may have benefits in reducing EC growth and arterial remodeling in PAH.

We also identified a novel, relatively de-dedifferentiated EC population in the PAH model, dDECs, that emerged early in disease and exhibited reduced expression of tight junction related genes such as *Cdhn5* and *Cldn5*, indicative of a loss of endothelial integrity, along with upregulation of antigen presentation related genes such as *Cd74* and *RT1-Da*. Interestingly, upregulation of CD74 expression was previously reported in the endothelium of pulmonary arteries and cultured ECs from PAH lungs.^60^ As a receptor for macrophage migratory inhibitory factor (MIF), CD74 may contribute to inflammatory cell recruitment which is well known to be associated with arterial remodeling in PAH.^36^ Enriched expression of *Cd74* was also described in a single cell transcriptomic analysis of lung ECs isolated from a SU/CH mouse model of PH;^12^ however, this was seen across multiple EC populations including capillary, arterial, and venous. As well, the mouse model, unlike the rat, does not exhibit occlusive arterial remodeling and the authors found no evidence of EC proliferation in their single cell transcriptomic analysis.^12^ In our study, we were able to determine that *Cd74* enriched ECs represented a discrete subset of ECs that were characterized by a relatively dedifferentiated state. As well, we demonstrated a clear transcriptomic signature for cell proliferation, which was represented in the UMAP as a ‘proliferative node’ consisting of cells from several clusters, including the aAECs and dDECs, both of which expanded markedly during PAH progression.

In addition to any role in promoting vascular inflammation, our study provides novel evidence that dDECs that are primed to undergo endothelial to mesenchymal transition (EndMT) to give rise to vascular stromal cells, a process that has been long thought to contribute to adventitial remodeling and arterial stiffness in PAH.^45,61^ EndMT shares many similarities with epithelial to mesenchymal transition (EMT),^46^ and is a phenomenon characterized loss of expression of canonical genes necessary for functional integrity, typically cell-cell junction proteins, with the adoption of motile and invasive cell behaviors. Interestingly, our RNA velocity analysis showed numerous strong vectors within the dDEC cluster directed towards various fibroblast populations. Moreover, latent time analysis showed that dDECs approximated a progenitor cell fate whereas the gCap EC populations were closer to a ‘terminal’ state of cell differentiation. While we were able to infer increased activity of TFs implicated in EndMT in this population, such as *Zeb2* and *Nr2f2*, it has increasingly been recognized that EndMT is a complex process and cannot be defined based on a small number of molecular markers.^46^ Indeed, dDECs showed a number of other features consistent with EndMT, including a marked reduction in the activity of *Erg*, a ‘master’ endothelial TF that promotes endothelial homeostasis via regulation of lineage-specific enhancers and super-enhancers.^47^ Indeed, a previous study showed that knockdown of *Erg* together with another ETS family transcription factor, *Fli1*, was sufficient to induce EndMT.^48^ Notably, the dDEC cluster was the only endothelial population that exhibited a very low activity of both *Erg* and *Fli1* (**Figure 8D**). Therefore, it is not surprising that canonical endothelial tight junction genes were downregulated in dDECs, along with increased activity of pathways associated with inflammation and apoptosis, which have been implicated in EndMT,^62,63^ as well as selective upregulation of several other EndMT related genes, including *Tpt1*,^37^ *Crip1*,^64^ and *S100a4*.^61^

Therefore, we used serial single cell transcriptomics to identify disease-specific EC and stromal cell populations that play a central role in arterial remodeling during the progression of PAH. Notably, we show that *Cd74*-enriched dDECs represent a distinct EC population exhibiting features of dedifferentiation and loss of functional integrity, that are primed to undergo EndMT, thereby contributing to the generation of profibrotic arterial fibroblasts that play an important in adventitial remodeling in PAH. As well, we have demonstrated that the aAEC cluster exhibits a marked growth dysregulated transcriptional state, possibly in response to ongoing hemodynamic perturbation, that disproportionally impacts the distal lung arteriolar bed. As well, for the first time, we provide evidence that the aAEC and SM-like pericyte populations are spatially localized to regions of complex arterial remodeling and contribute to intimal obliteration and distal muscularization, respectively, in both the SU/CH model and PAH patients. Moreover, since they are characterized by high *Tm4sf1* expression, this may be a promising target for therapeutic strategies to prevent or reverse arterial remodeling in PAH.

## Acknowledgements

We would like to thank Anli Yang for her technical assistance with all animal experiments and Elmira Safaie Qamsari for her assistant sectioning fresh frozen tissues for immunofluorescent staining. Additionally, we thank the following Ottawa Hospital Research Institute core services Stemcore laboratories for single cell library preparation and sequencing, OHRI Bioinformatics core for preprocessing of single cell data, and the University of Ottawa’s Animal Care and Veterinary Services core.

## Author Contributions

NDC has contributed to all aspects of this work including conception, experimental design, performing the experiments, data analysis and preparation of this manuscript. EM contributed to data analysis and figure preparation. RSG assisted with collection of the first experiment of single cell samples and preliminary analysis. YD was instrumental in preparation and collection of the lung samples for microCT analysis and in all animal studies. KS performed the IPA analysis. AS performed the RNA velocity analysis and contributed to figure preparation. DPC assisted with single cell transcriptomic analysis and provided valuable support in initial experimental design. SEL performed immunofluorescent staining on all human samples and contributed to figure preparation. TK performed immunofluorescent staining on fresh frozen rat sections. KY collected images on rat lung sections, assisted with figure preparation and data interpretation, and manuscript review. SB provided access to human PAH samples, contributed expertise to interpretation of results, and manuscript review. DJS was involved in the conception, experimental design, data analysis and interpretation, figure design, and drafting the manuscript. DJS is accountable for all aspects of this work.

