## Supplemental Figures + Legends for "Emergence of disease-specific endothelial and stromal cell populations responsible for arterial remodeling during development of pulmonary arterial hypertension"

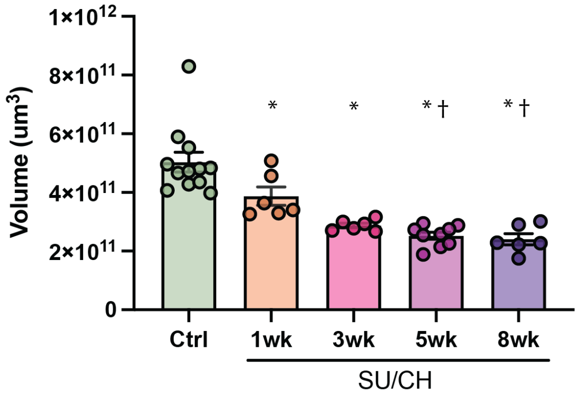


**Supplemental Figure 1: Loss of pulmonary vascular volume during SU/CH.** Total vascular volume demonstrating progressive loss throughout SU/CH development. Data represented as mean ± SEM, n = 6 – 12 biological replicates, * $p < 0.05$ vs healthy control, † $p < 0.05$ vs 1-week.

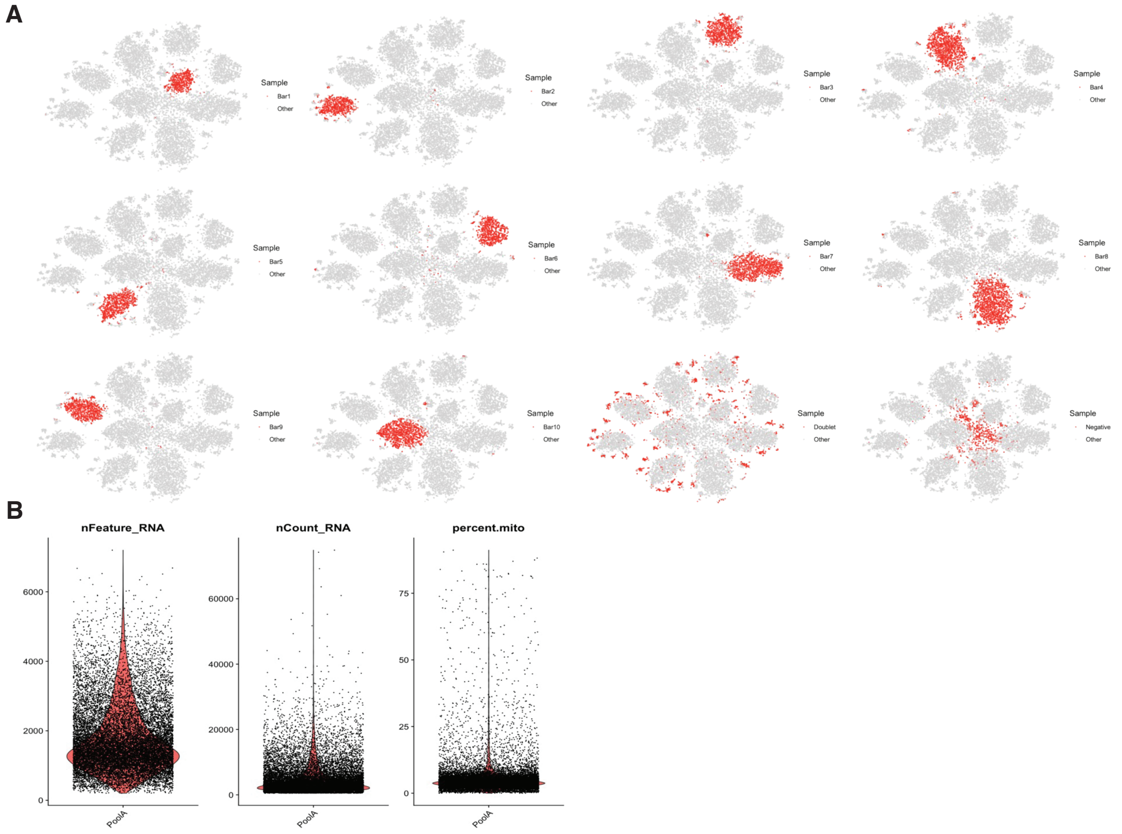


**Supplemental Figure 2: Quality control metrics during deMultiplex and preprocessing.** (A) Representative plots from one lane of 10x Genomics samples while preforming demultiplexing. Cells strongly expressing individual barcodes clustered together (singlets), cells expressing multiple barcodes are seen in small clusters on the periphery (doublets), and cells with insufficient barcode expression clustered in the middle (negative). Only singled identified barcodes were kept. (B) Additional quality control preprocessing in Seurat using features, counts, and mitochondrial content.

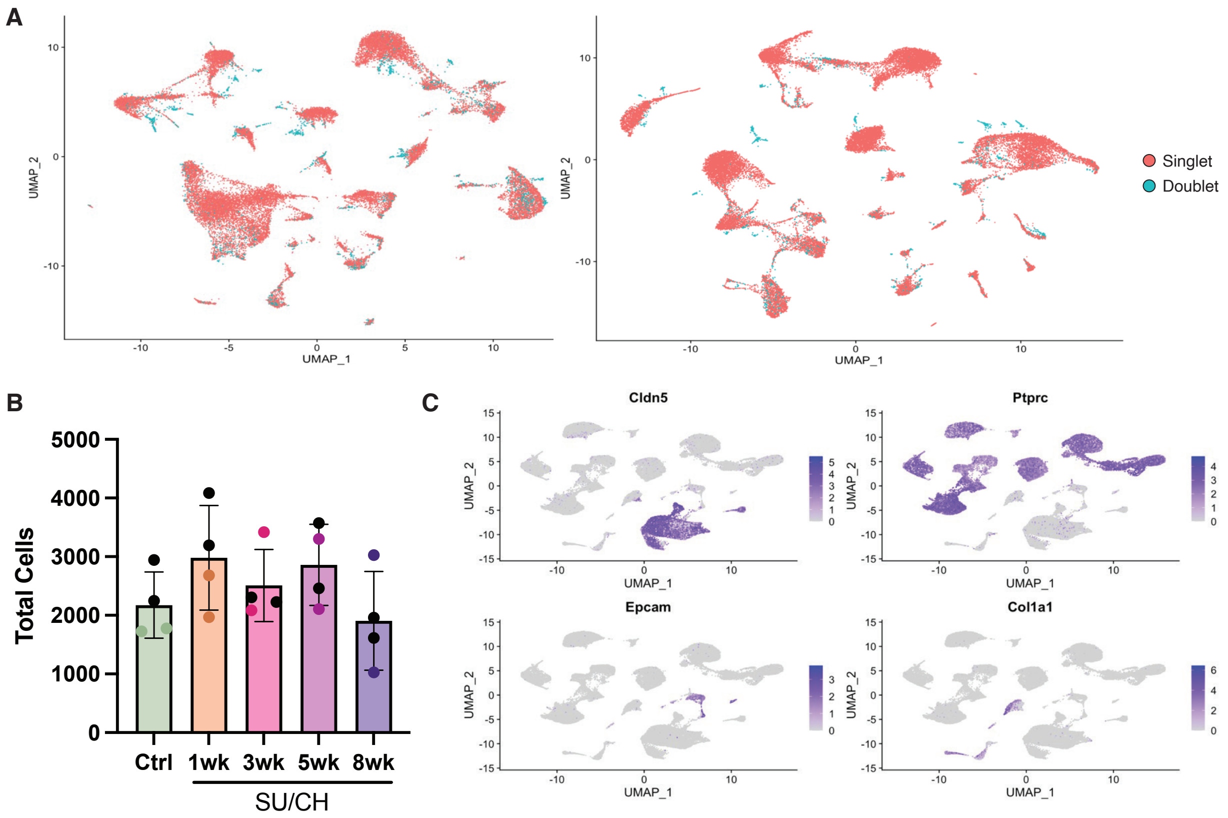


**Supplemental Figure 3: Doublet removal and bulk population identification.** (A) Prior to merging and integrating the two experiment an additional round of doublet removal was performed with scDblFinder. (B) Total cells for each biological sample in the integrated Seurat object. (C) Identification of major lung cell populations based on common bulk identify genes *Cldn5* (endothelial), *Ptprc* (immune), *Epcam* (epithelial), and *Col1a1* (stromal).

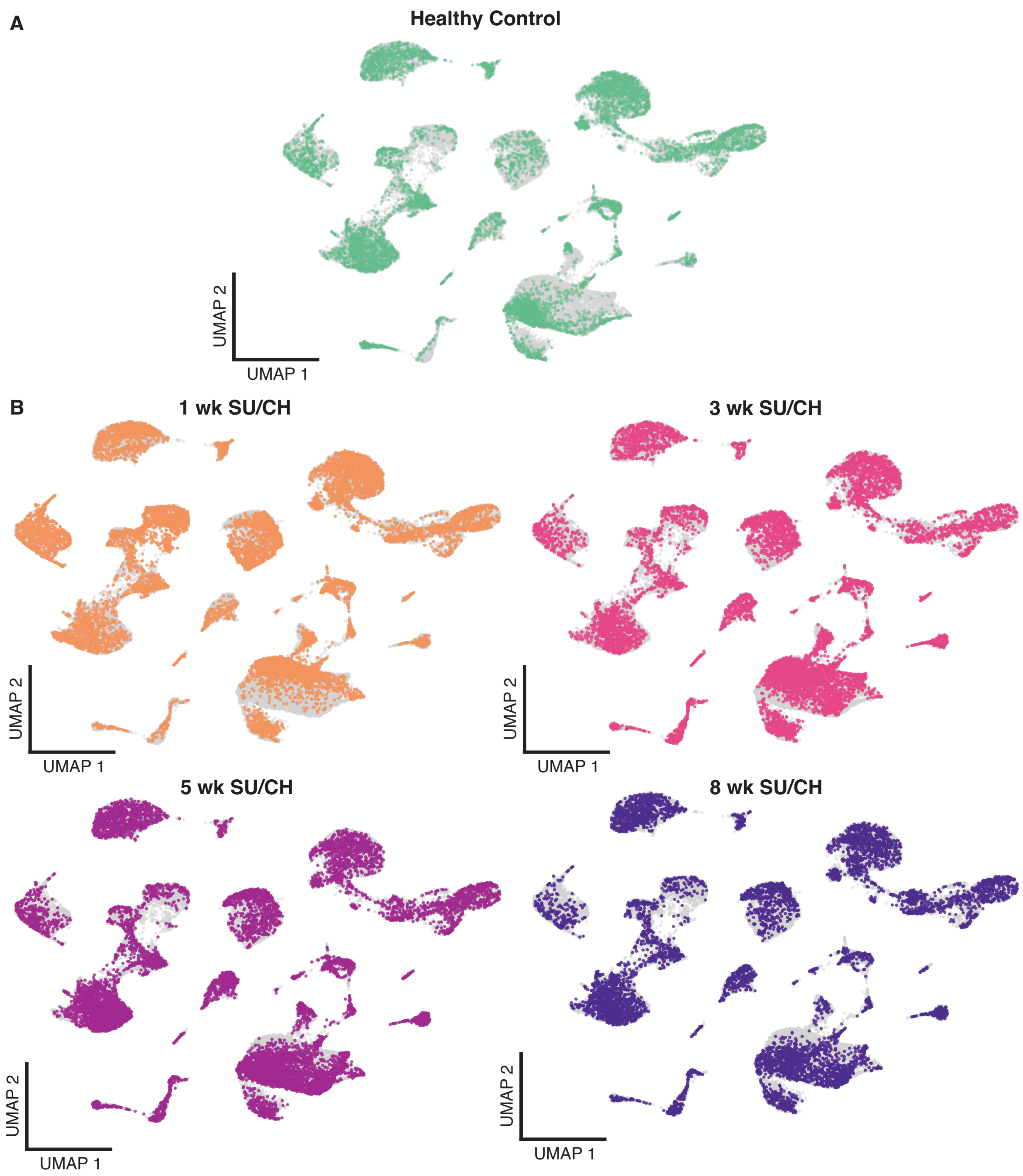


**Supplemental Figure 4: Global UMAPs by timepoint.** Representative UMAPs showing each timepoint control – green (A), 1wk SU/CH – orange (B), 3 wk SU/CH – pink (C), 5 wk SU/CH – purple, 8 wk SU/CH – dark purple.

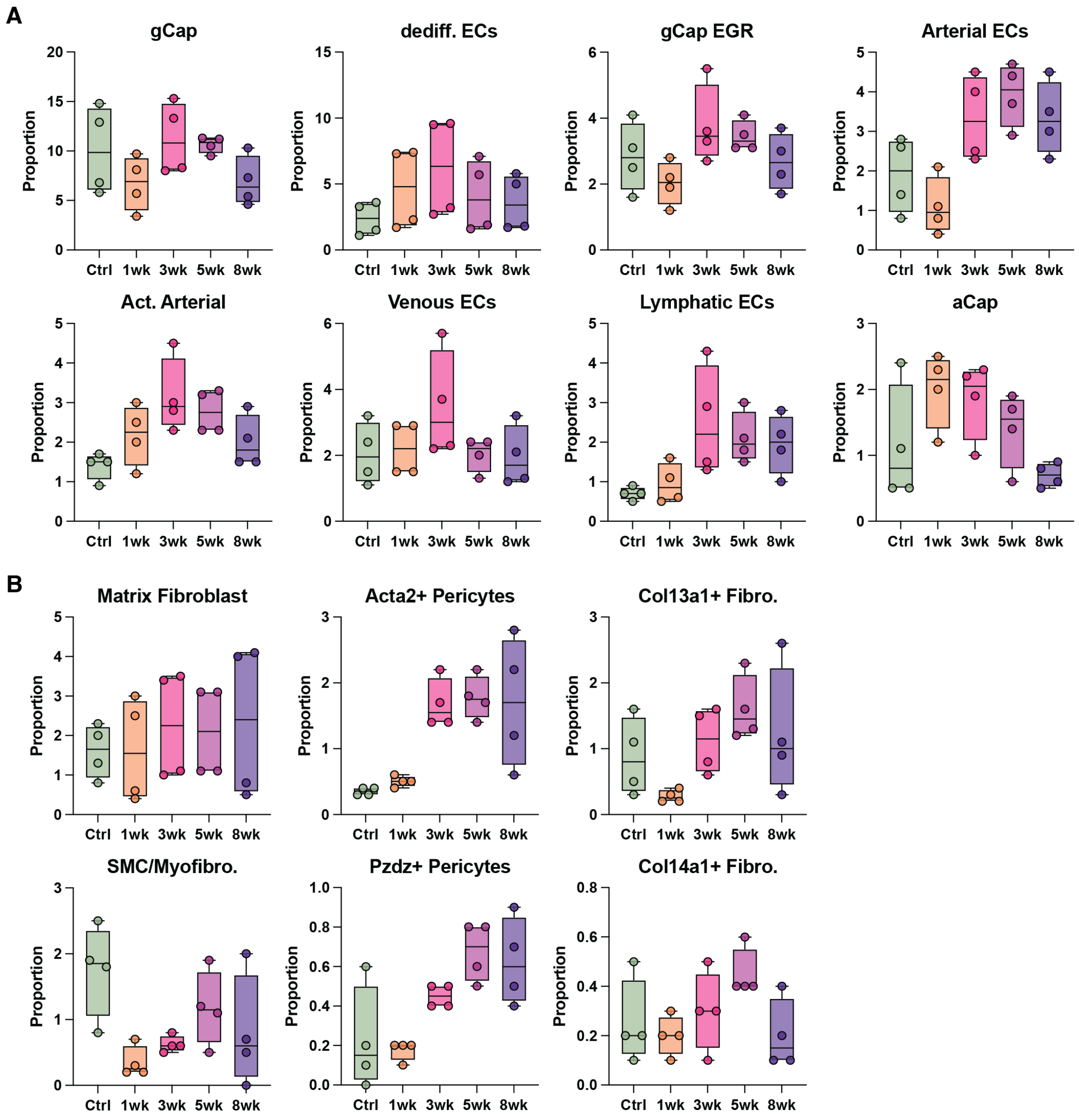


**Supplemental Figure 5: Global proportions of all endothelial and cells within integrated Seurat object.** Changes in global proportions of cell populations were quantified in the endothelial (A) and stromal (B) subsets by timepoint, demonstrating relative increases in arterial ECs, activated arterial ECs, *Acta2*+ pericytes, and classical pericytes during PAH progression compared to controls. Representing n = 4 biological replicates per timepoint.

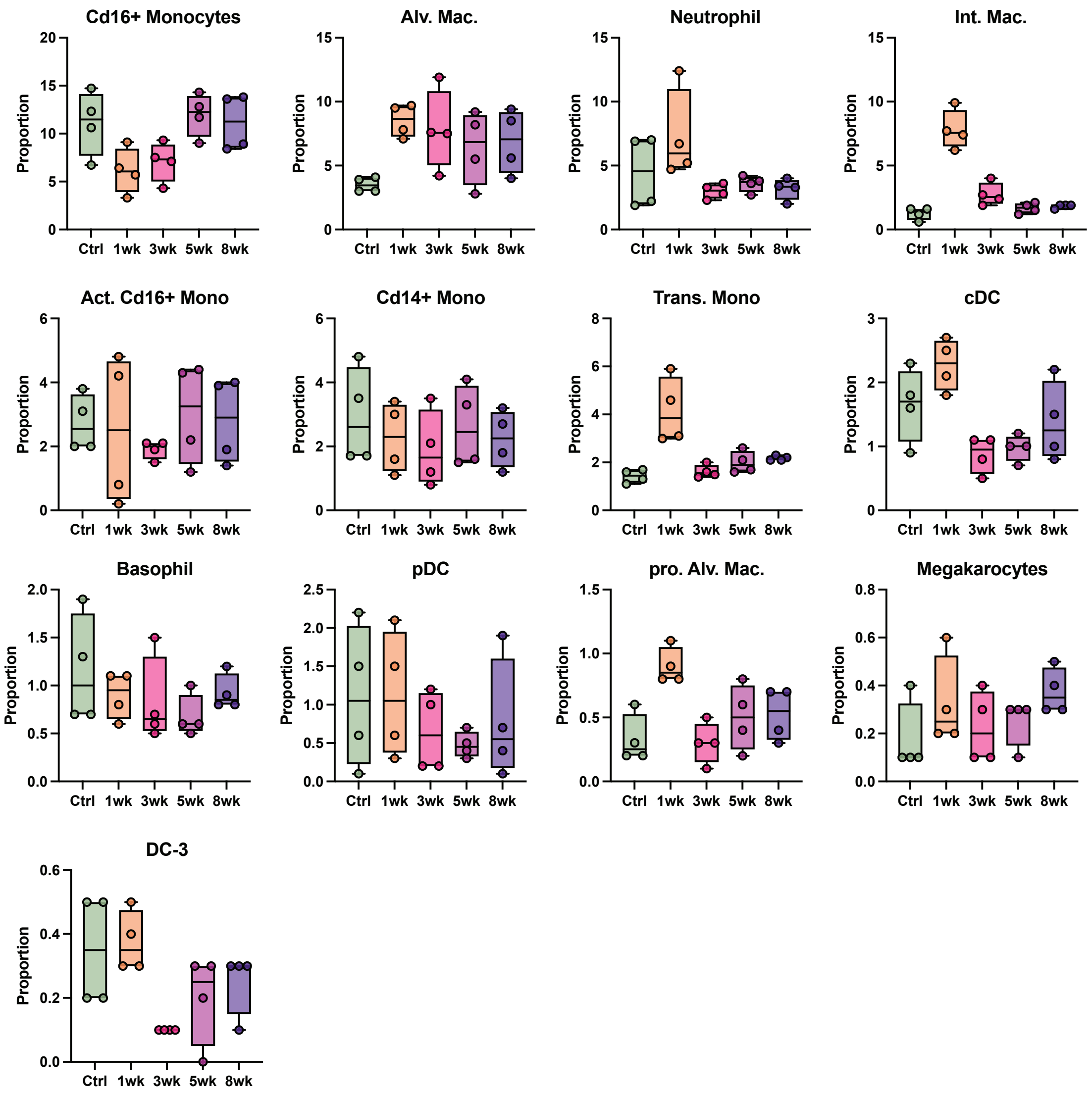


**Supplemental Figure 6: Global proportions of all myeloid cells within integrated Seurat object.** Representing n = 4 biological replicates per timepoint.

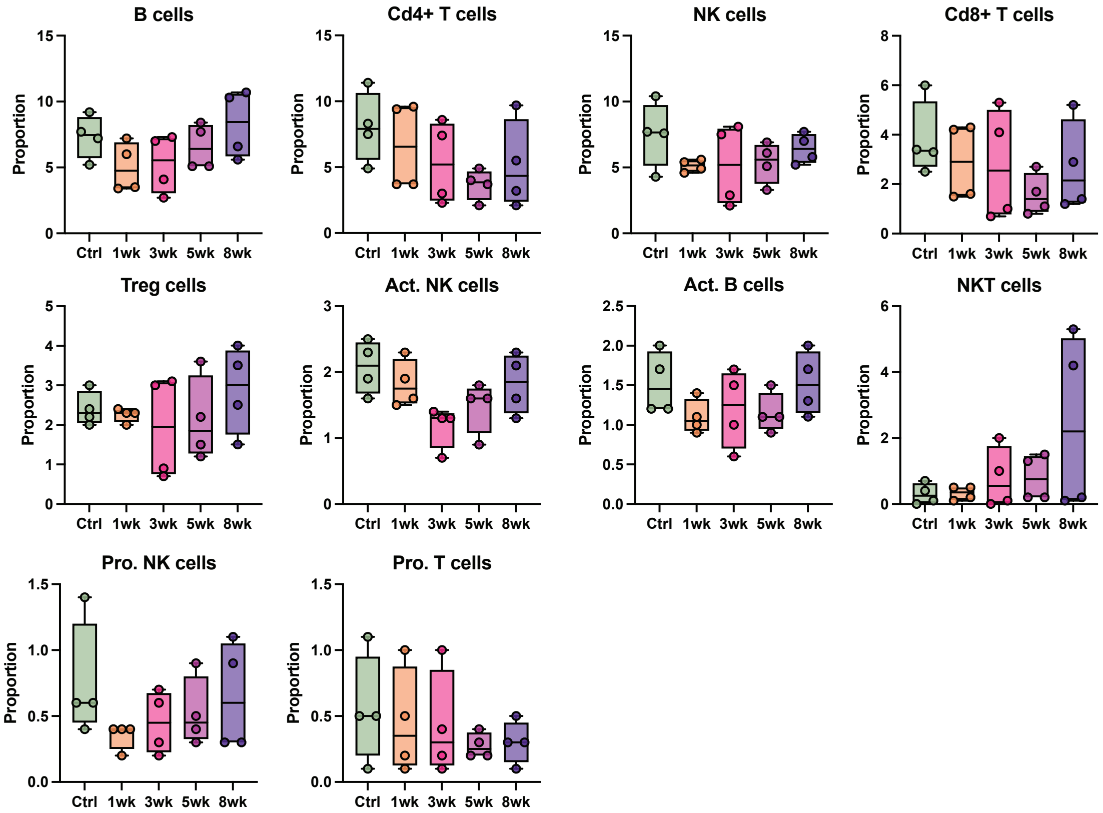


**Supplemental Figure 7: Global proportions of all lymphoid cells within integrated Seurat object.** Representing n = 4 biological replicates per timepoint.

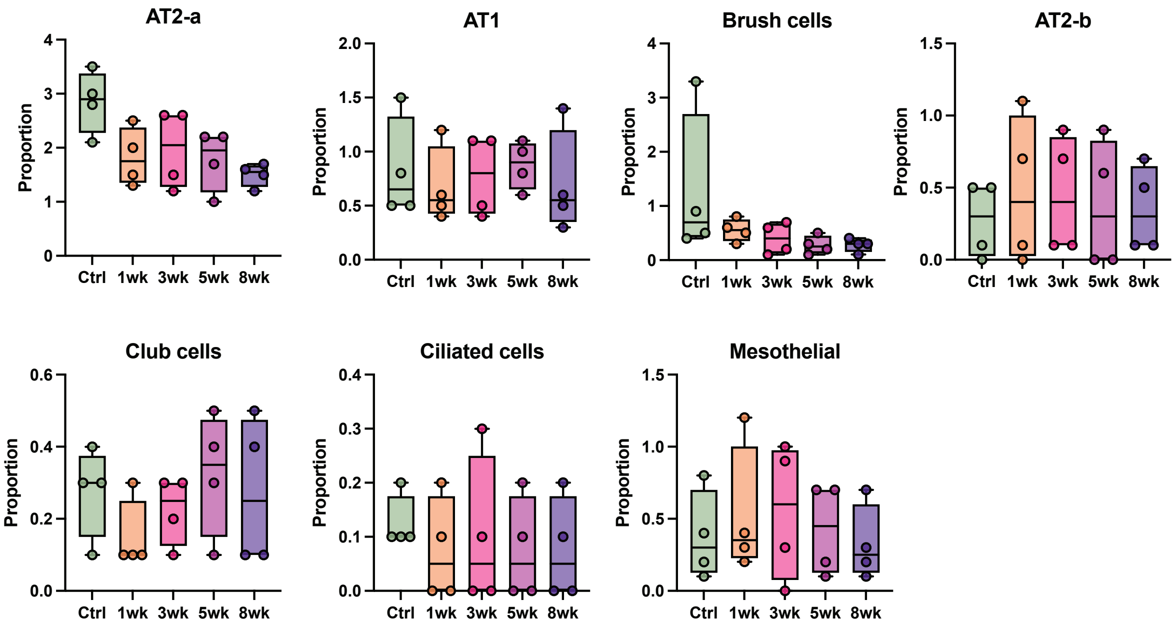


**Supplemental Figure 8: Global proportions of all epithelial and mesothelial cells within integrated Seurat object.** Representing n = 4 biological replicates per timepoint.

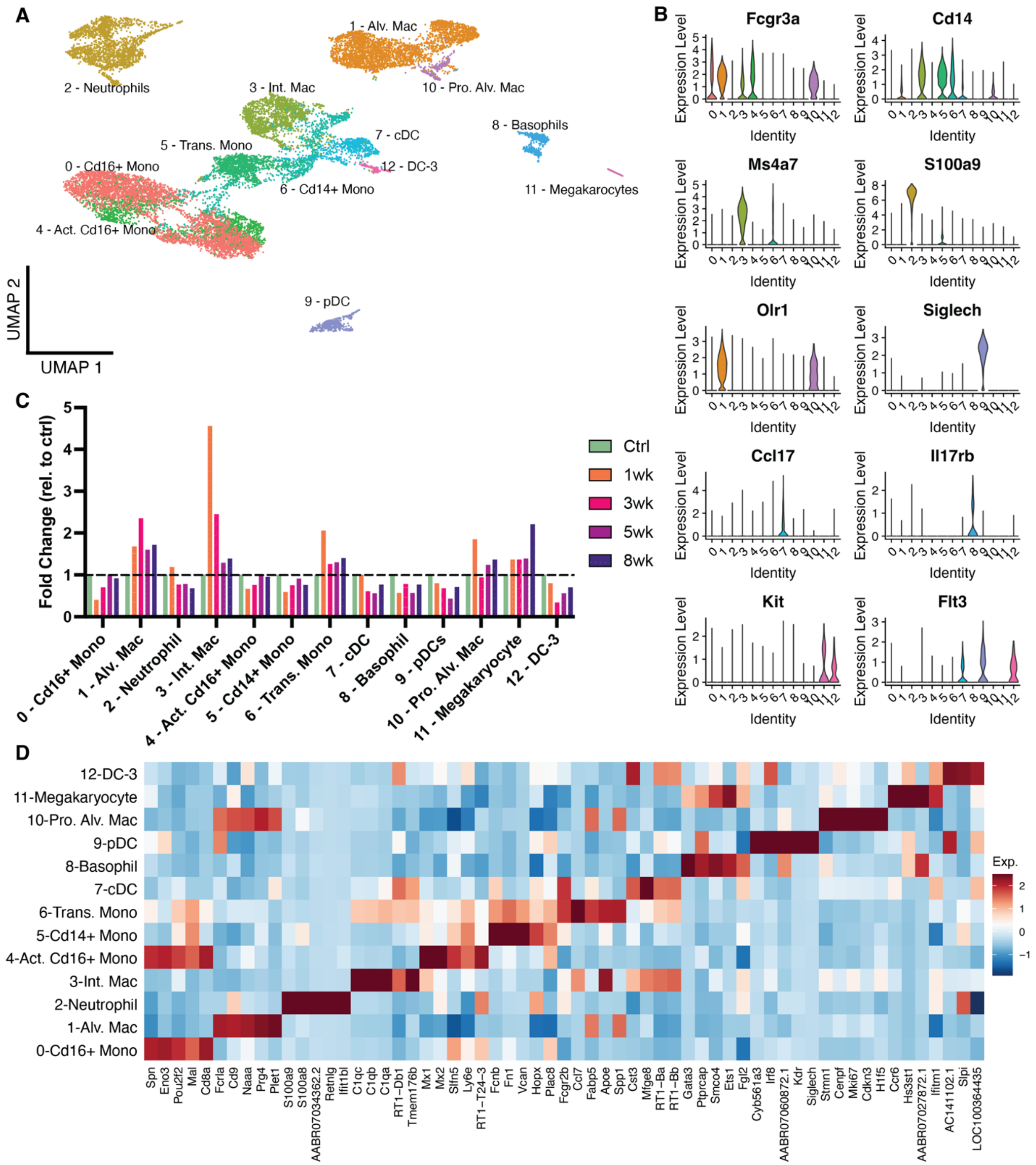


**Supplemental Figure 9: Myeloid subcluster analysis.** (A) UMAP of myeloid subset representing 13 resolved cell types. (B) Top genes used to identify myeloid populations. (C) Fold change of myeloid populations at each timepoint relative to healthy control samples. (D) Heatmap of the top 5 differentially expressed gene per myeloid cluster.

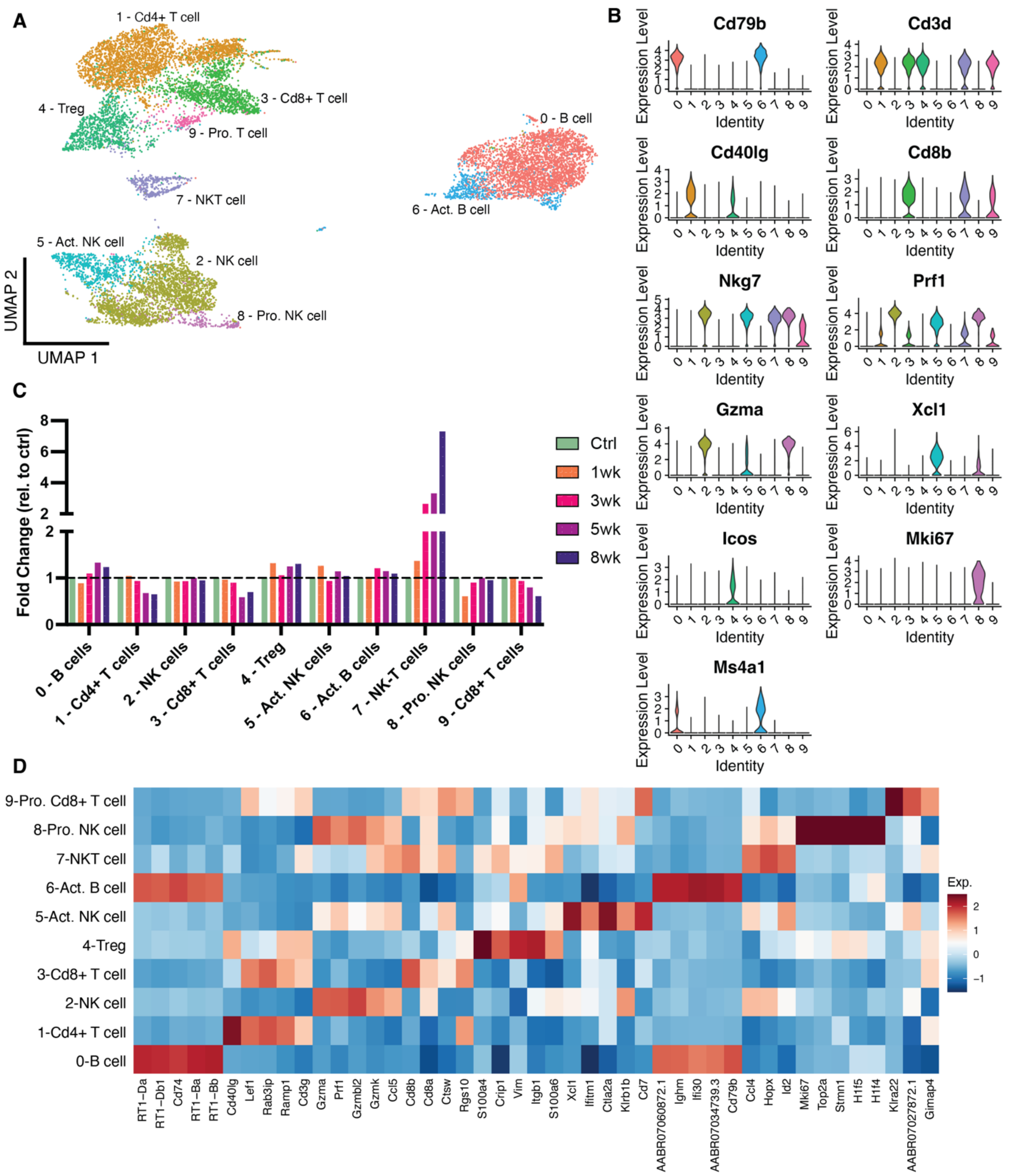


**Supplemental Figure 10: Lymphoid subcluster analysis.** (A) UMAP of lymphoid subset representing 10 resolved cell types. (B) Top genes used to identify lymphoid populations. (C) Fold change of lymphoid populations at each timepoint relative to healthy control samples. (D) Heatmap of the top 5 differentially expressed gene per lymphoid cluster.

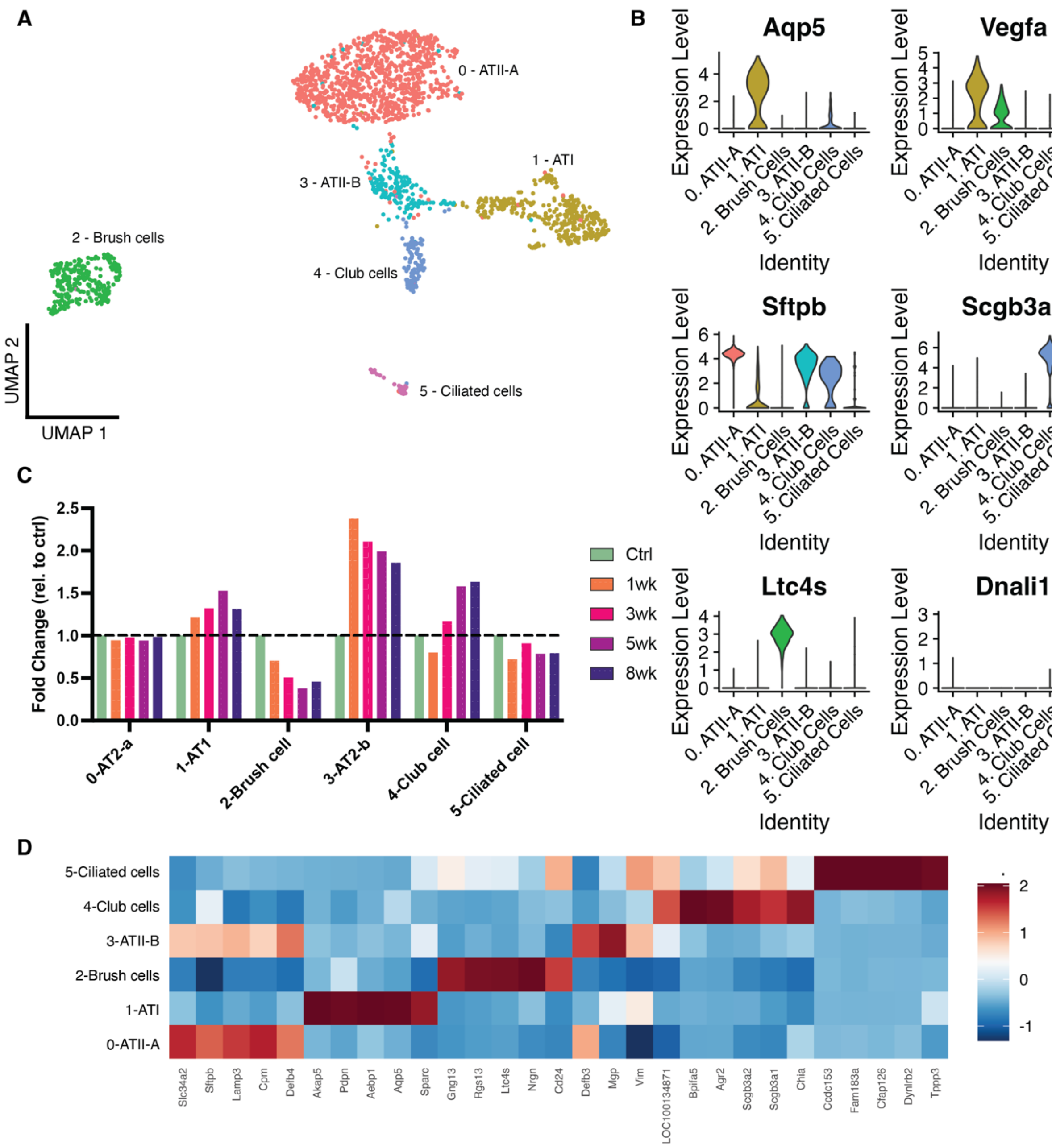


**Supplemental Figure 11: Epithelial subcluster analysis.** (A) UMAP of epithelial subset representing 6 resolved cell types. (B) Top genes used to identify epithelial populations. (C) Fold change of epithelial populations at each timepoint relative to healthy control samples. (D) Heatmap of the top 5 differentially expressed gene per epithelial cluster.
